# A dual-clock-driven model emulating the effects of either Ano1 or IP_3_R knock-out on lymphatic muscle cell pace-making

**DOI:** 10.1101/2022.11.06.515373

**Authors:** E.J. Hancock, S.D. Zawieja, C. Macaskill, M.J. Davis, C.D. Bertram

## Abstract

Lymphoedema, a common dysfunction of the lymphatic system, results in fluid accumulating between cells. Fluid return through the lymphatic vascular system is primarily provided by contractions of muscle cells in the walls of lymphatic vessels, driven by electrochemical oscillations causing rhythmic action potentials and associated surges in intracellular calcium ion concentration. There is incomplete understanding of the mechanisms involved in these repeated events, restricting the development of pharmacological treatments for dysfunction. Previously, we proposed a model where autonomous oscillations in the membrane potential (M-clock) drove passive oscillations in the calcium concentration (C-clock). In this paper, to model more accurately what is known about the underlying physiology, we extend this model to the case where the M-clock and the C-clock oscillators are both active but coupled together, and thus both driving the action potentials. This extension results from modifications to the model for the IP_3_ receptor, a key C-clock mechanism. The synchronized dual-driving clock behaviour enables the model to match IP_3_ receptor knock-out data, resolving an issue with previous models. We also use phase-plane analysis to explain the mechanisms for the dual-clock coupling. The model has the potential to help determine mechanisms and find targets for pharmacological treatment of lymphoedema.

## 1 INTRODUCTION

The lymphatic system vasculature is composed of a network of vessels, nodes and valves that returns interstitial fluid from between cells back to the blood circulation, and operates in parallel with the blood circulatory system (Moore & Bertram 2018). It also plays a vital role in the immune system, as the returning fluid is filtered via lymph nodes for a range of particles, bacteria, and viruses. Lymphatic system defects are involved in numerous diseases, such as obesity, cardiovascular disease, inflammation, and neurological disorders such as Alzheimer’s disease (Mortimer & Rockson 2014, Oliver et al. 2020).

Lymphoedema is a long-term dysfunction of the lymphatic system wherein the fluid is not properly returned to the circulatory system and instead accumulates in the interstitium (Rockson 2001). It consists of sustained swelling in the affected region, and often results in secondary infections (Mortimer & Rockson 2014) and significant reduction in quality of life (Jørgensen et al. 2021, Zhang et al. 2021). The dysfunction is commonly acquired secondary to events such as lymphatic filariasis, congestive heart failure and cancer treatment. It affects over 10 million people annually in the USA (Rockson & Rivera 2008) and over 130 million people worldwide (Mortimer & Rockson 2014). While decongestive therapy and surgical interventions can help treat the symptoms, and even improve lymphatic function (Adams et al. 2010), there is currently no treatment of the underlying causes.

The return of interstitial fluid via the lymphatic vascular system is driven by both intrinsic and extrinsic pumping, where intrinsic pumping results from periodic contractions of the lymphatic muscle in the vessel walls, and accounts for about two-thirds of lymph flow in the extremities at rest (Olszewski & Engeset 1980). These contractions are presumed driven by the autonomous pace-making ability of lymphatic muscle cells. The remainder of the fluid flow is due to extrinsic pumping, which results from intermittent passive squeezing of surrounding tissues. Both pumping mechanisms depend on competent one-way lymphatic valves (Breslin et al. 2019). Overall, lack of intrinsic lymph pumping is central to lymphoedema, and consequently it is important to study the pace-making ability of lymphatic muscle cells (LMCs) that drives this pumping.

However, there is incomplete understanding of the mechanisms involved in the oscillatory excitation for intrinsic pumping, restricting the development of pharmacological treatments (Lee et al. 2022). Due to the complexity of these pace-making systems, modelling is critical to uncovering the function and failure of the underlying oscillatory mechanisms. While LMCs are almost uniquely complex^1^ in mounting both tone and short-lived contractions (Muthuchamy et al. 2003), for the inbuilt pacemaker behind the latter they can draw extensively from models of oscillations in other cell types. Models of oscillatory mechanisms in a range of cell types have historically been based on either a so-called ‘membrane clock’ (M-clock) as originated by Hodgkin & Huxley (1952) or a ‘calcium clock’ (C-clock); see, e.g., Goldbeter et al. (1990). The M-clock is composed of periodic flows of ion currents in the cell membrane, resulting in an oscillating membrane voltage output consisting of rhythmic action potentials separated by a period of relative quiescence. Propagation of the action potential along the fibre as a signal is the purpose in nerve as studied by Hodgkin & Huxley, and in the conduction system of the heart. The C-clock represents the oscillating levels of calcium in the cell, and models are composed of the calcium concentrations in the main part of the cell (cytosol), the calcium store (endoplasmic reticulum, ER) and mathematical descriptions of the calcium flux across the ER and cell membranes. In muscle, where the store is termed the sarcoplasmic reticulum, the periodic peaking^2^ of [Ca^2+^], giving rise to mechanical contraction, is the purpose. Models of pace-making in cardiac cells were based around the M-clock (McAllister et al. 1975), while a range of other cell types have focussed on oscillations in C-clocks (Dupont et al. 2016). More recently there has been a shift, backed by experimental data, in the models of cardiac pace-making to dual coupled clocks, where neither clock is dominant in triggering action potentials (Yaniv et al. 2015).

Lymphatic muscle has so far seen limited modelling of pacemaker mechanisms. Hald et al. (2018) created a two-variable model of a voltage oscillator (i.e. M-clock only), based on modifications to a barnacle muscle model (Morris & Lecar 1981). This model was created in order to study communications between neighbouring LMCs and had limited detail for internal mechanisms. Imtiaz et al. (2007) added M-clock components to an earlier calcium-based model by Dupont & Goldbeter (1993). In this model the C-clock drives the oscillations and triggers the action potential in the M-clock. In previous work, we extended the model of Imtiaz et al. by modifying mechanisms in the M-clock (Hancock et al. 2022): specifically, by altering the model components for the L-type calcium channel. These modifications enabled the M-clock to oscillate independently and drive oscillations in the C-clock, which was necessary for the model to match experimental data based on knock-out of anoctamin-1 (Ano1) calcium-activated chloride-ion channels. These channels form a key M-clock mechanism enabling coupling from the C-clock.

In this paper we direct attention to the inositol 1,4,5-triphosphate (IP_3_) receptor, which is known (Hille 2001, Emrich et al. 2021) to be the main source of calcium ion release from the store to the cytoplasm in smooth muscle types, including lymphatic muscle. As such it is essential for the LMC’s C-clock, and its numerical knock-out automatically stops pace-making in any model such as that of Imtiaz et al. (2007) which is C-clock driven. However, this is not the case experimentally. Figure 1 shows a comparison of action potentials (APs) recorded from mouse LMCs under matched control conditions and after deletion of the dominant IP_3_ receptor 1 (IP_3_R1-KO)^3^. Clearly APs survive, albeit with reduced frequency. The systolic plateau phase is lengthened after IP_3_R-KO, and the total minimum-to-maximum excursion of the membrane potential is less.

**Figure 1.**
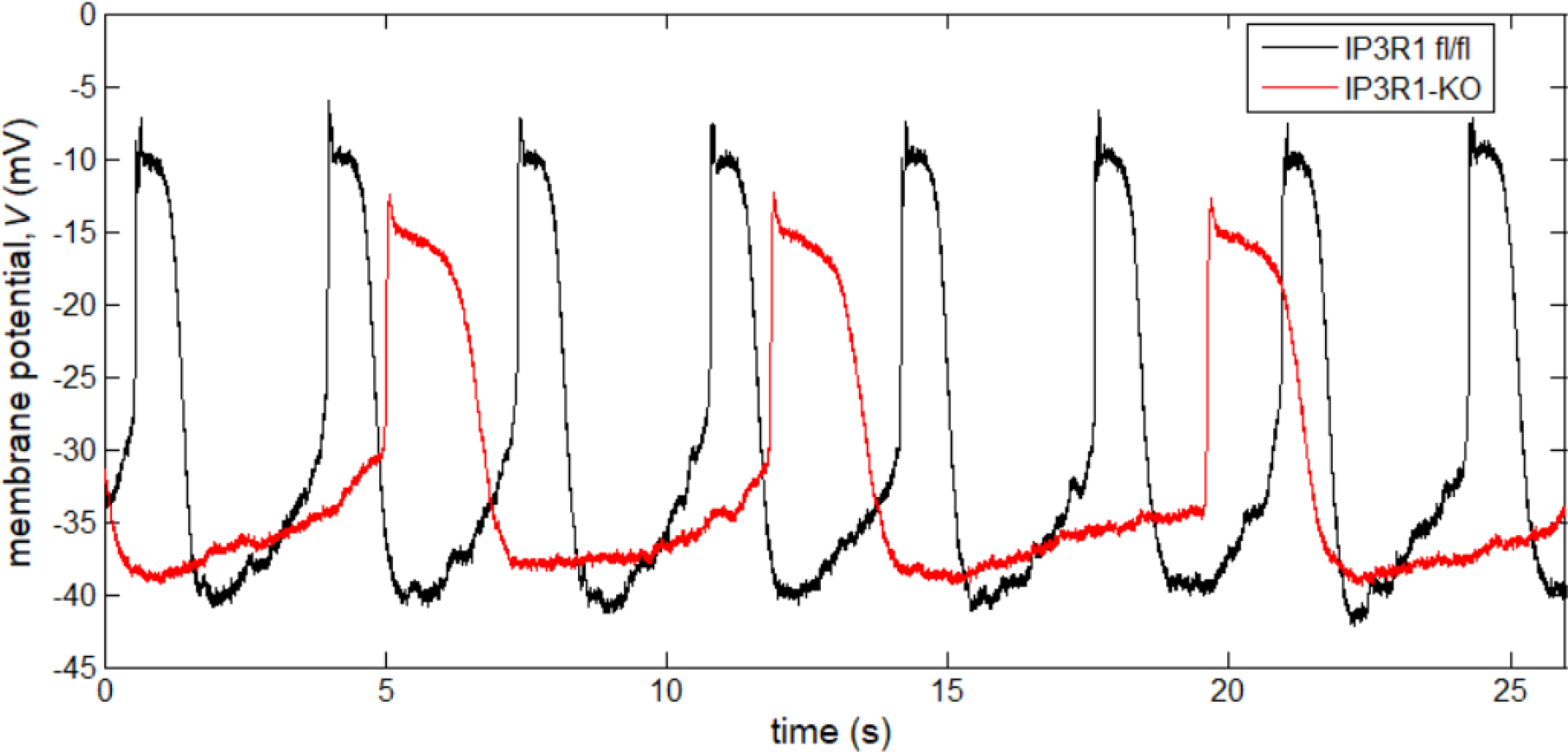
Membrane potential vs. time, recorded from murine LMCs *ex vivo*. Here, the frequency of APs averages 17.8 /min under control conditions (black trace), and 8.2 /min under IP_3_R-KO (red trace).

The extent of the frequency reduction after IP_3_R-KO is variable; Figure 2 shows another example. Whereas in Fig. 1 the frequency reduces to 46% of control, in Fig. 2 it reduces to less than 37% of control^4^. Again the systolic plateau phase is pronouncedly lengthened after IP_3_R-KO, and the total excursion is less. In Fig. 2 the total excursion is greater under control conditions than in Fig. 1, because of both a lower minimum immediately after repolarization and a higher maximum early-systolic spike, whereas the IP_3_R-KO waveform in Fig. 2 displays a similar excursion to that in Fig. 1, but the whole waveform is raised relative to its Fig. 1 counterpart, so that the membrane potential is considerably higher than control during the diastolic slow depolarization phase.

**Figure 2.**
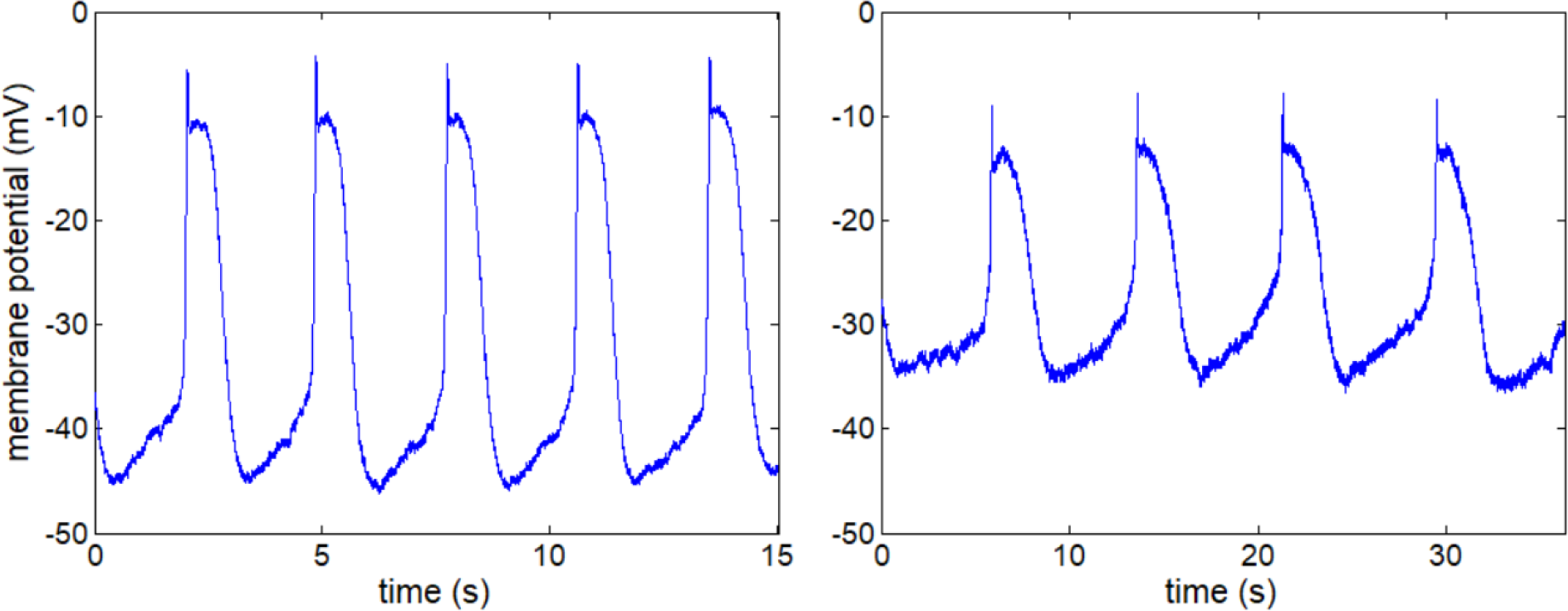
Membrane potential vs. time, recorded from murine LMCs *ex vivo*. Left panel: wild-type control. Right panel: IP_3_R1-KO (note different time-base). Here, the AP frequency averages 20.9 /min under control conditions and 7.6 /min under IP_3_R-KO.

In this paper, by modifying the C-clock mechanisms, we extend our existing model to the case where the M-clock and the C-clock are both driving oscillations. We first analyse our existing model to show how that model fails to emulate the observations of knock-out of the IP_3_ receptor. We then analyse a special case of the proposed model to illustrate two different means by which the C-clock can control oscillation frequency in the M-clock: the first, by varying the frequency of C-clock oscillations, and the second, by varying the [Ca^2+^] level of a non-oscillating C-clock. Finally, we analyse the proposed model for the general case and show that the model largely matches with features of the IP_3_R-KO experimental data. We use time-traces and phase-plane analysis to show that the C-clock and the M-clock are both driving their synchronised oscillations.

The rest of the paper is set out as follows. Section 2 provides background on the existing model that we extend in this paper. Section 3 describes the new model and methodology used in this paper. Section 4 presents the results from the proposed model, including the case where both clocks are driving oscillations and phase-plane analysis explanations. Section 5 provides a discussion of the results. Section 6 draws brief conclusions.

## 2 BACKGROUND

### 2.1 Existing model and problem with IP3R-KO simulation

In this section we provide a background on our existing model [see Hancock et al. (2022) for details]. That model matched the experimentally observed membrane potential oscillations under both control and Ano1-KO conditions, with Ano1-KO leading to larger action potentials (APs) and a much longer period, as seen in Figure 3. However, Fig. 3 also shows the membrane potential computed with the IP_3_ receptor disabled (IP_3_R-KO) by setting the maximum IP_3_R flux *J*_x3_ to zero [see table of notation and equations of Hancock et al. (2022)]. The added traces show that, under IP_3_R-KO, once start-up transients decay, the existing model produces APs which are similar in all respects to those produced under control conditions. This is not consistent with the observed IP_3_R-KO behaviour as shown in Figs. 1 and 2. The data for this latter numerical case were not part of the original study and are a focus of the work in the current paper.

**Figure 3.**
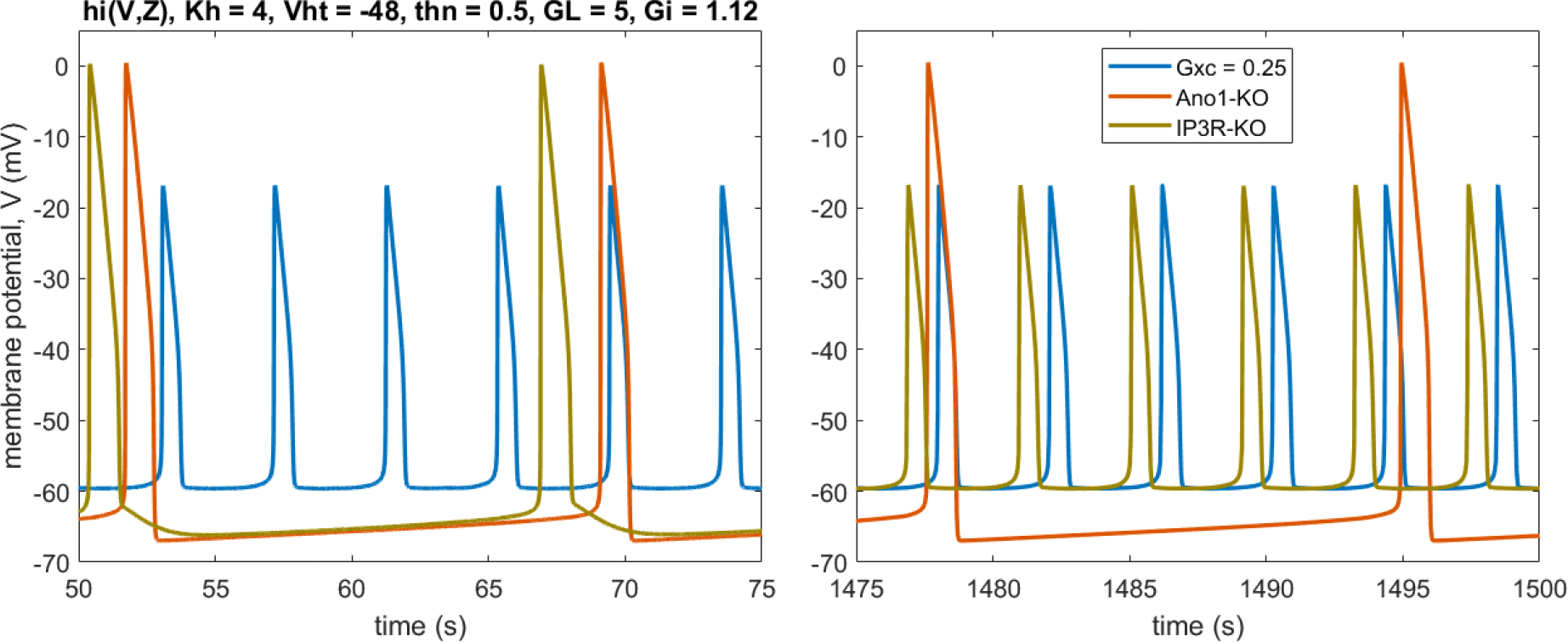
Time-traces from existing model by Hancock et al. (2022) with traces for IP_3_R-KO added. Note large change in IP_3_R-KO waveform size and frequency between early and late simulated time, owing to gradual increase of store [Ca^2+^] to an asymptotic value.

The existing model is composed of an M-clock and a C-clock using four time variables. The C-clock is represented by two calcium variables: the cytosolic calcium concentration *Z*, and the endoplasmic reticulum (ER) stored calcium concentration *Y*. The M-clock is represented by two membrane variables: the membrane potential *V* and an inactivation gating variable *h* for the L-type calcium channel; see eqs. 1–4 and fig. 2 of Hancock et al. (2022).

Hancock et al. (2022) used phase plane analysis to analyse the oscillations of the clocks. Phase-plane analysis shows the relationship between two different time-variables on a single graph of their combined trajectory, rather than plotting them individually against time; see Strogatz (2015) for further details. To simplify analysis, the phase-plane analysis for the two clocks was separated, as shown in Figure 4. The trajectories of the two clocks were plotted on their respective phase planes. The nullclines of the M-clock were also plotted, whilst assuming that the (C-clock-dependent) calcium concentration *Z* was constant – see Fig. 4. Similarly, the nullclines of the C-clock were plotted assuming that (M-clock-dependent) L-type calcium current *I_L_* was held constant. A nullcline of a variable shows all points where that variable does not change with respect to time. It is useful for interpreting the phase plane, since it divides the plane into regions where the time derivative of the variable has different signs.

**Figure 4.**
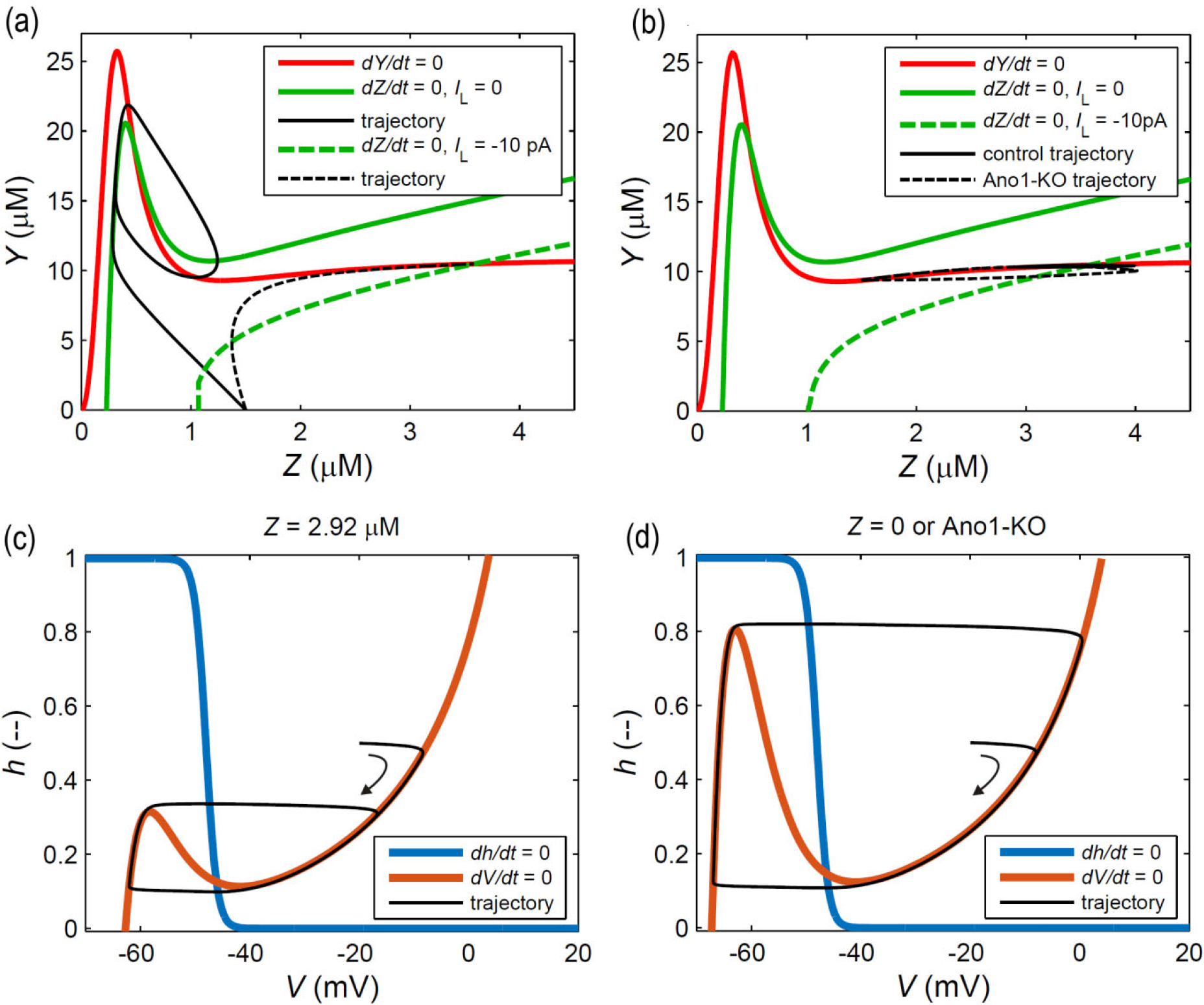
Phase planes for existing model by Hancock et al. (2022). (a) Phase plane for C-clock isolated by fixing *I*_L_. (b) C-clock phase plane with full clock coupling. (c) Phase plane for M-clock isolated by fixing *Z*. (d) As 4c but with zero *Z* or, equivalently, Ano1-KO. With *K*_h_ = 4 μM as in Fig. 3, the *h*-nullcline is invariant over the *Z*-range 0 to 2.92 μM.

In the existing model the M-clock drove oscillations of both [Ca^2+^] and membrane potential. This was required in order to match the Ano1-KO data. In the model of Imtiaz et al. (2007), the M-clock was driven by oscillations arising in the C-clock. Under these circumstances oscillations of membrane potential (APs) were impossible with Ano1-KO, because the latter condition removed the mechanism for the C-clock to drive the M-clock. In Fig. 4a, increasing *Z* depresses the peak of the *V*-nullcline. Consequently the M-clock orbital trajectory is relatively small, indicating increased M-clock frequency and decreased peak membrane potential *V*. This is exemplified in Fig. 4 by moving from panel (d) to panel (c). Conversely, Ano1-KO, which for the M-clock amounts to *Z* = 0, leads to the maximum sized orbit, indicating decreased AP frequency accompanied by increased peak *V* and lowered minimum *V*.

In the existing model the C-clock had the means to oscillate independently when uncoupled from the M-clock, but this did not occur when the M-clock was connected. In Fig. 4a & 4b the *Z*-nullcline is shown for two fixed values of *I_L_*, the first (*I_L_* = 0) pertaining to when the C-clock was disconnected from the M-clock, and the second (*I_L_* = −10 pA) exemplifying values occurring when the clocks were coupled. Fig. 4a shows trajectories from an arbitrary starting location for the C-clock independent of the M-clock. It can be observed that the C-clock oscillated between relatively low values of *Z* when *I_L_* = 0 but approached a stable equilibrium at a considerably higher value of *Z* when *I_L_* = −10 pA. Fig. 4b shows the orbital trajectories when the C-clock was coupled to the M-clock. In Fig. 4b, *Z* oscillates between extremes of 2.9 and 4.0 μM under control (WT) conditions (solid black line), and between 1.5 and 4.0 μM with Ano1-KO (dashed black line). Under both conditions, the orbit surrounds a nullcline intersection point that is stable (Fig. 4a), showing that the C-clock is not here oscillating in its own right. For this case, the oscillations are driven by changes in *I_L_* and thus by the M-clock.

### 2.2 Modification of the existing model

In this paper, we focus on modifying and analysing the C-clock model, as modelling IP_3_R mechanisms are central to fitting the IP_3_R-KO data. To achieve this, we focus on two questions. First, can the C-clock model be improved by using more modern mechanistic approaches [see Dupont et al. (2016) for background on calcium modelling]? Second, can the C-clock and M-clock each drive oscillations in the other, rather than just the M-clock driving the C-clock?

## 3 METHODS

### 3.1 Experiments

A moderately detailed description of the biological methods was given by Hancock et al. (2022); the companion paper to this one (Zawieja et al., submitted) gives full details, including the genetic and pharmacological means whereby IP_3_R knock-out was secured.

### 3.2 Description of the new model

In this section we introduce a model for LMC oscillations by improving on our previous model of LMC pace-making (Hancock et al. 2022). We modify the model of the C-clock to alter the coupling between the M- and C-clocks and the effect of IP_3_. This modification incorporates a previous IP_3_R model by De Young & Keizer (1992), as simplified by Li & Rinzel (1994).

The model assumes a single well-mixed space constituting the cytoplasm, and a single internal compartment, the sarcoplasmic reticulum (SR); see Figure 5. It incorporates both the M-clock and C-clock oscillations using five ordinary differential equations (ODEs), and thus five interacting time-variables. The C-clock is represented by three variables: the cytosolic calcium concentration *Z*, the SR calcium concentration *Y,* and a gating variable *n* that modulates the time-course of the calcium release from the SR. The M-clock is represented by two membrane variables: the membrane potential *V* and an inactivation gating variable *h* for the L-type calcium channel. All other variables are algebraically related to one or more of these five. The C-clock consists of the calcium-channel fluxes through the cell membrane and between the cytosol and the SR. The M-clock relates the membrane potential to all the currents through the membrane. The model ODEs are

**Figure 5.**
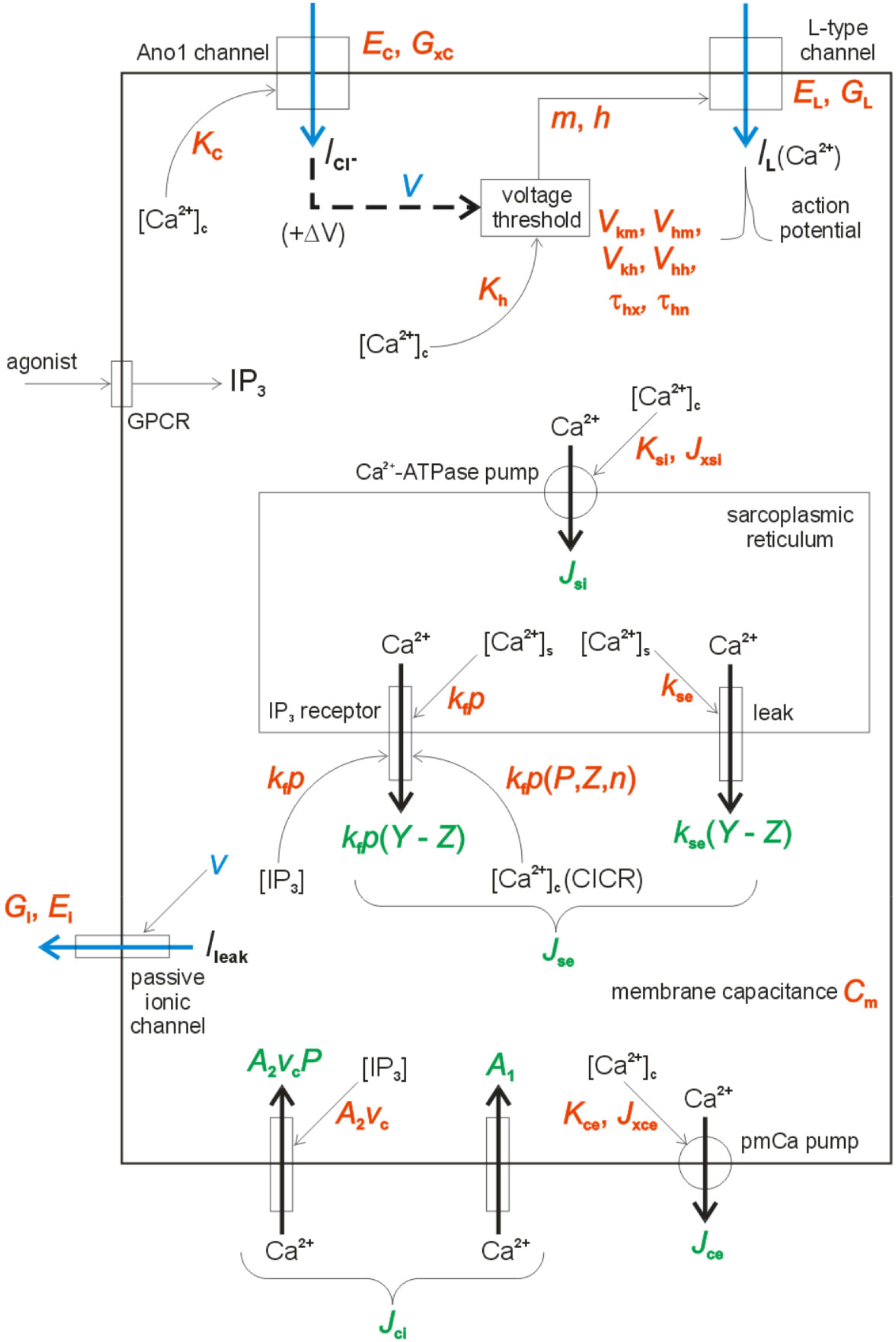
Schematic of the new LMC pacemaker model. Currents are shown by blue arrows, and calcium fluxes by black arrows. By convention, currents indicate the direction that positive charges, real or imaginary, are moving; thus an outward current of negatively charged chloride ions is here shown as an inwardly directed current of fictitious positive charges, producing an increase in (trans)membrane potential (defined as internal relative to external).

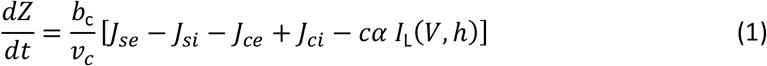

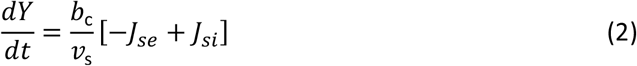

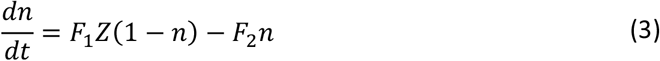

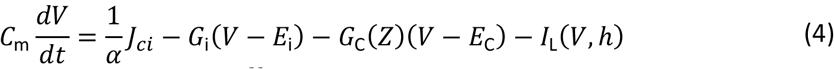

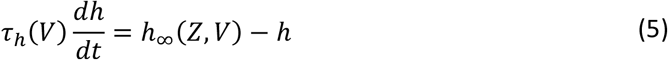

where *J*_ci_ is the Ca^2+^ influx across the membrane into the cell from outside, *J*_ce_ represents the pumping of Ca^2+^ from the cytosol out of the cell*, J*_si_ is the Ca^2+^ influx from the cytosol into the store, and *J*_se_ is the release of calcium from the store into the cytosol. *P* is the cytosolic concentration of IP_3_ (herein chosen to remain constant), and *F*_1_ and *F*_2_ are functions of *P*. For the membrane voltage, *C*_m_ represents the capacitance of the cell membrane, *G*_i_ and *E*_i_ represent a lumped conductance and equilibrium potential respectively for passive non-selective ionic channels, *G*_C_ and *E*_C_ represent a lumped conductance and equilibrium potential respectively for the Ano1 channels, and *I_L_* represents the summed current in the L-type calcium channels^5^. *h_∞_* and *τ_h_* represent the steady state and voltage-dependent time constant respectively for the L-type inactivation gating variable *h*. *v*_c_ is the volume of the cytoplasm, *v*_s_ is the volume of the SR, *b*_c_ represents the fraction of total calcium that is unbuffered, and *α* is a conversion factor between current and calcium ion flux. These functions are described by

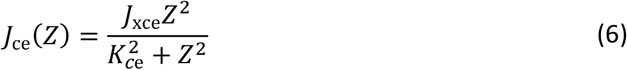

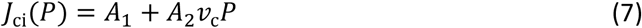

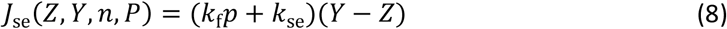

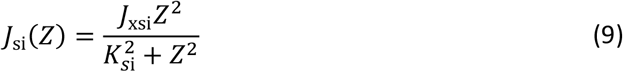

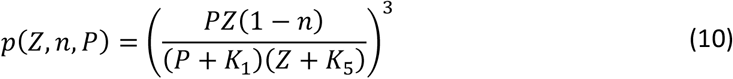

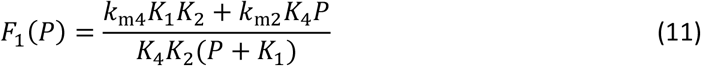

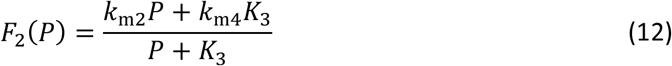

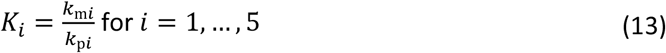

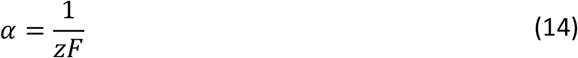

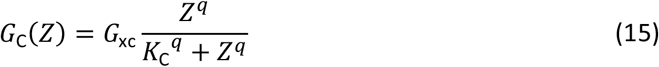

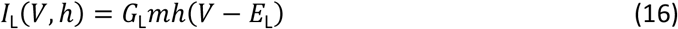

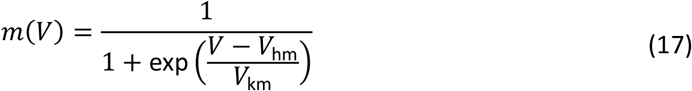

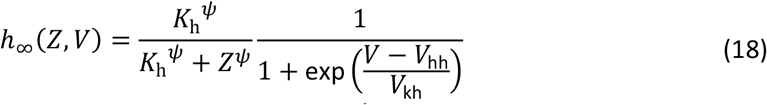

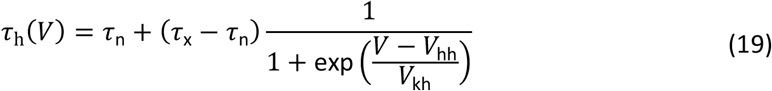

where the values of all constants are given in Table 1. The L-type current is controlled by the activating and inactivating gate variables, *m* and *h* respectively, where *m* is assumed to act instantaneously and *h* is subject to first-order delay kinetics according to the time constant *τ_h_*, itself now ranging between maximum and minimum values *τ_x_* and *τ_n_* respectively according to the membrane potential.

In the newly adopted IP_3_R model, calcium release from the store is a function of store [Ca^2+^] and cytosolic [IP_3_] and [Ca^2+^], the latter property representing calcium-induced calcium release (CICR). While the former IP_3_R model included less exact representations of all these, the new model, via the time-dependent properties imbued by the gating variable *n*, also emulates the sequential activation and inactivation of the IP_3_R (Dupont et al. 2016).

**Table 1.** Values for constants in the model. Sources (otherwise, estimated): 1 is Imtiaz et al. (2007), although in many cases the values have been reinterpreted in terms of units, etc., 2 is Lees-Green et al. (2014), 3 is Keener & Sneyd (2009), and 4 is a universal physical constant. Estimated values were based on existing sources.

| parameter | units | meaning | value | source |
| --- | --- | --- | --- | --- |
| $J_{xce}$ | $\mu\text{mol/s}$ | max. $\text{Ca}^{2+}$ efflux from cell via pmCa pump | $0.28 v_c/b_c$ | |
| $K_{ce}$ | $\mu\text{M}$ | half saturation Z-value for pmCa pump | 0.425 | |
| $A_1$ | $\mu\text{mol/s}$ | constant $\text{Ca}^{2+}$ leak into cell | $0.0015 v_c/b_c$ | |
| $A_2$ | /s | $\text{Ca}^{2+}$ flux into cell per unit of $\text{IP}_3$ | $0.01/b_c$ | |
| $P$ | $\mu\text{M}$ | $\text{IP}_3$ concentration in cytoplasm | 0.7 | |
| $J_{xsi}$ | $\mu\text{mol/s}$ | maximum SERCA flux | $9 v_c/b_c$ | |
| $K_{si}$ | $\mu\text{M}$ | half saturation Z-value for SERCA pump | 0.1 | |
| $k_{se}$ | L/s | constant $\text{Ca}^{2+}$ leak from store | $3.3 v_c/b_c$ | |
| $k_f$ | L/s | scale for $\text{IP}_3$-dependent store $\text{Ca}^{2+}$ efflux | $4.8 v_c/b_c$ | |
| $k_{p1}$ | /$\mu\text{mol/s}$ | $\text{IP}_3\text{R}$ binding constant | 400 | 3 |
| $k_{p2}$ | /$\mu\text{mol/s}$ | $\text{IP}_3\text{R}$ binding constant | 0.2 | 3 |
| $k_{p3}$ | /$\mu\text{mol/s}$ | $\text{IP}_3\text{R}$ binding constant | 400 | 3 |
| $k_{p4}$ | /$\mu\text{mol/s}$ | $\text{IP}_3\text{R}$ binding constant | 0.2 | 3 |
| $k_{p5}$ | /$\mu\text{mol/s}$ | $\text{IP}_3\text{R}$ binding constant | 20 | 3 |
| $k_{m1}$ | /s | $\text{IP}_3\text{R}$ binding constant | 52 | 3 |
| $k_{m2}$ | /s | $\text{IP}_3\text{R}$ binding constant | 0.21 | 3 |
| $k_{m3}$ | /s | $\text{IP}_3\text{R}$ binding constant | 377.2 | 3 |
| $k_{m4}$ | /s | $\text{IP}_3\text{R}$ binding constant | 0.029 | 3 |
| $k_{m5}$ | /s | $\text{IP}_3\text{R}$ binding constant | 1.64 | 3 |
| $b_c$ | - | $\text{Ca}^{2+}$ buffering constant | 0.01 | 2 |
| $v_c$ | L | cytosol volume | $10^{-12}$ | 2 |
| $v_s$ | L | store volume | $v_c/5.5$ | 3 |
| $C_m$ | pF | cell capacitance | 25 | 2 |
| $E_i$ | mV | reversal potential, passive channels | -67.2 | 1 |
| $E_C$ | mV | reversal potential for Ano1 current | -20 | 1 |
| $E_L$ | mV | reversal potential for L-type channels | +20 | 1 |
| $G_i$ | nS | conductance of passive channels | 0.4 | |
| $G_{xC}$ | nS | max. $Ca^{2+}$-induced $Cl^-$ conductance | 0.8 | |
| $G_L$ | nS | conductance of L-type $Ca^{2+}$ channels | 1.7 | |
| $V_{hm}$ | mV | half saturation for gate fn. $m(V)$ | -46 | |
| $V_{km}$ | mV | inverse slope const., gate fn. $m(V)$ | -4 | |
| $K_h$ | $\mu M$ | half saturation for gate fn. $h_\infty(Z)$ | 4 | |
| $\psi$ | - | Hill exponent for $h_\infty(Z)$ | 20 | 1 |
| $V_{hh}$ | mV | half saturation for gate fn. $h_\infty(V)$ | -48 | |
| $V_{kh}$ | mV | inverse slope const., gate fn. $h_\infty(V)$ | 1 | |
| $\tau_n$ | s | minimum of time constant $\tau_h(V)$ | 0.5 | |
| $\tau_x$ | s | maximum of time constant $\tau_h(V)$ | 10 | |
| $\alpha$ | ($\mu mol/s$)/pA | $Ca^{2+}$ flux per unit current | $5.18 \times 10^{-12}$ | 4 |
| $z$ | - | valency of calcium ion | 2 | 4 |
| $F$ | pC/$\mu mol$ | Faraday constant | $9.6485 \times 10^{10}$ | 4 |

A finite-difference version of the above equations was coded in Matlab and solved on a single processor of a standard PC. Nullclines were found, in most cases iteratively, by solving the appropriate equation from eqs. 1 to 5 with the time derivative set to zero. All results shown are from times much later than required for start-up transients to decay, apart from those for the existing model in the left panel of Fig. 3.

### 3.3 Nullcline intersection stability

For stability analysis of a phase plane, we can represent a two-dimensional system in a general form by

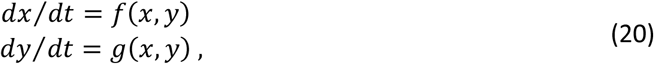

where *x* and *y* are the variables (e.g. *V* and *h* for the M-clock here) and *f* and *g* are the functions representing the changes in *x* and *y* with respect to time. Note that *f* is here associated with the abscissa variable, and *g* with the ordinate. At equilibrium we have *f*(*x*_0_, *y*_0_) = 0 and *g*(*x*_0_, *y*_0_) = 0, i.e. (*x*_0_, *y*_0_) is a point in (*V*, *h*)-space where neither *V* nor *h* is changing with time. Since the two nullclines of a phase plane are curves along which *f* = 0 for one and *g* = 0 for the other, their intersection point provides such an equilibrium point. However, what happens next depends on whether this equilibrium is stable or unstable. The equilibrium point (*x*_0_, *y*_0_) is stable if [see chs. 5 and 6 of Strogatz (2015) for details]

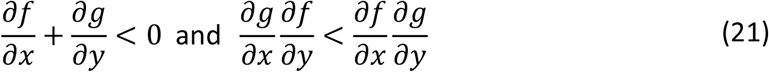

where the partial derivatives are determined at the values *x* = *x*_0_, *y* = *y*_0_. The equilibrium point is unstable if either of the inequality signs is reversed, meaning that any slight perturbation causes departure from (*x*_0_, *y*_0_).

The above analysis applies to the 2-dimensional M-clock of this paper, i.e. it can be used to determine whether an intersection between the nullclines *dV/dt* = 0 and *dh/dt* = 0 is stable or not. However, the C-clock is 3-dimensional, so in general the analysis cannot be applied. We portray the C-clock herein by way of 2-dimensional phase-plane slices of the (*Z*,*Y*,*n*)-space. Provided a single such slice, e.g. a phase plane in (*Z*,*n*)-space with *Y* = const., is considered, the analysis can be applied to the nullcline intersection(s) found therein.

## 4 RESULTS

### 4.1 New model

In this section we present time-traces and phase-plane analysis for the proposed models of oscillations in membrane potential and calcium concentration. We show different methods whereby the C-clock can control the oscillations of the M-clock. We also show that the proposed model qualitatively matches the experimental control, Ano1-KO and IP_3_R-KO data, and analyse the coupling of the M-clock with the C-clock.

#### 4.1.1 C-clock driving M-clock

We first analyse the two different ways that the C-clock can bring about action potential (AP) initiation in the M-clock. To analyse this triggering, we mathematically remove the effect of the M-clock on the C-clock while leaving the C-clock effect on the M-clock. We set the dimensionless binary constant *c* = 0 in eq. 1 above, which removes the effect of the L-type calcium channel on the cellular calcium concentration [Ca^2+^] while keeping its effect on voltage. This modification is equivalent to the L-type calcium channel passing not calcium but instead another bivalent cation, that changes the membrane potential *V* but not the cytosolic [Ca^2+^] (algebraically, *Z*). This modification allows us to study the temporal dynamics by which the M-clock is coupled to the C-clock.

We investigate two cases where the C-clock can modify the oscillations of the M-clock. In the first (*c* = 0), which will be the subject of Figures 6 and 7, the isolated C-clock can be at a quiescent steady state. In the second, which will be the subject of Figures 8–10, the isolated C-clock can spontaneously oscillate, and thereby manipulate the M-clock. The first case is idealised, as it could not occur for the real case of *c* = 1 if an action potential occurs. However, this case is informative as an approximation and a conceptual illustration. Figure 6 shows superimposed time-traces from the simulated system with *c* = 0, in the three situations of (1) control conditions (all parameters take their default values), (2) Ano1 channel knock-out (zero Ano1 conductance), and (3) IP_3_ receptor knock-out (zero Ca^2+^ efflux from the store via the IP_3_ receptor). The case of IP_3_R-KO displays constant [Ca^2+^], i.e. *Z*, after start-up transients decay, whereas the control case exhibits time-varying *Z*. Ano1-KO also displays time-varying *Z*, but will be regarded as a reference case in the sense that it is equivalent to *Z* = 0 as input to the Ano1 channel, thus disabling the mechanism by which the C-clock triggers APs in the M-clock.

**Figure 6.**
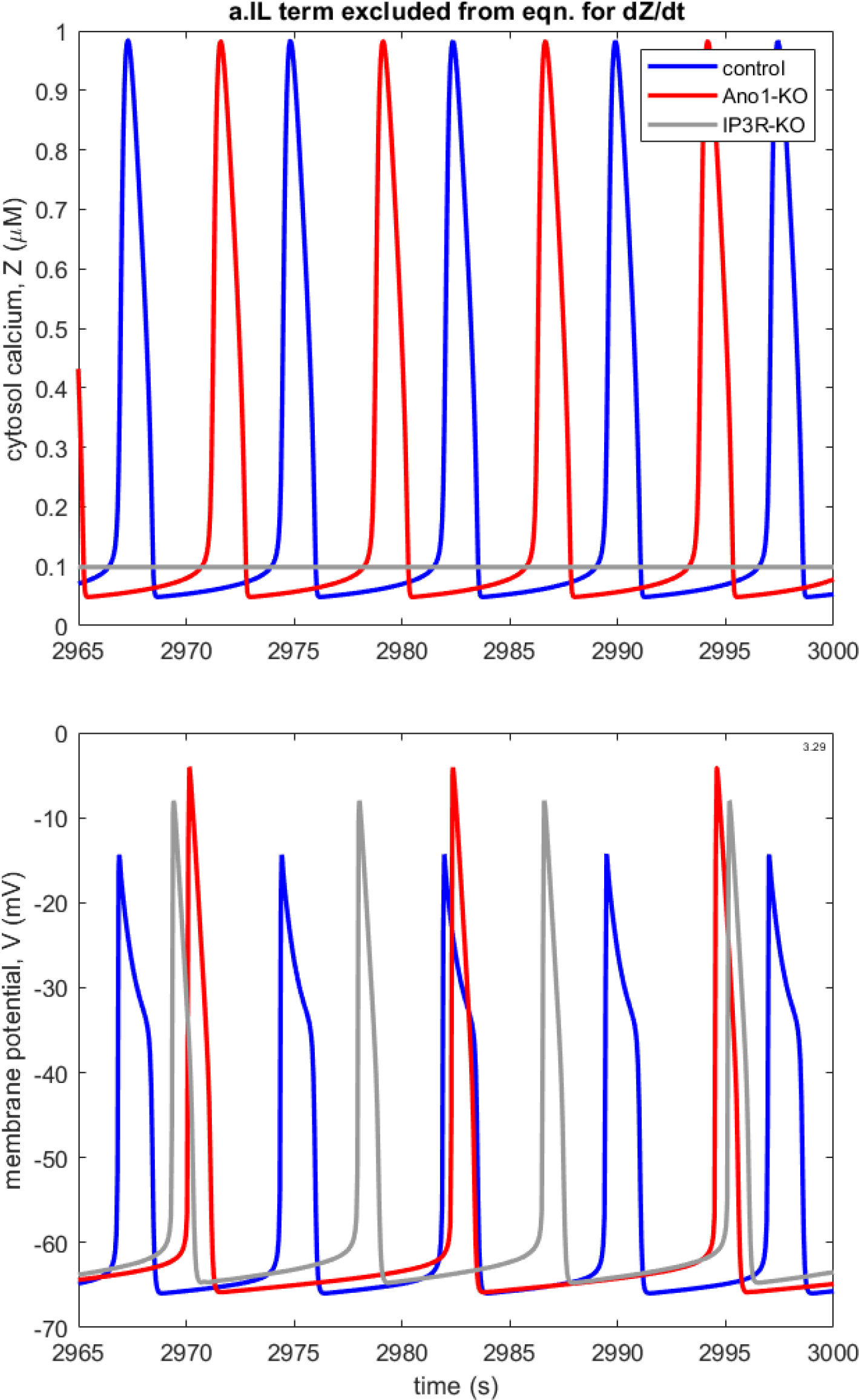
Superimposed traces from three runs with *c* = 0, showing waveforms after start-up transients have decayed, under control (blue), Ano1-KO (red) and IP_3_R-KO (grey). Top: cytosolic calcium concentration. Bottom: membrane potential. The AP frequencies are 8.0 (control), 4.9 (Ano1-KO) and 6.9 (IP_3_R-KO) /min. Parameter values that differ from those in Table 1: *P* = 0.9 μM, *J*_xce_ = 0.056*v*_c_/*b*_c_, *K*_ce_ = 0.1275 μM, *A*_1_ = 0.003*v*_c_/*b*_c_ μmol/s, *A*_2_ = 0.02/*b*_c_ /s, *K*_c_ = 0.3 μM, *V*_kh_ = 0.4 mV.

**Figure 7.**
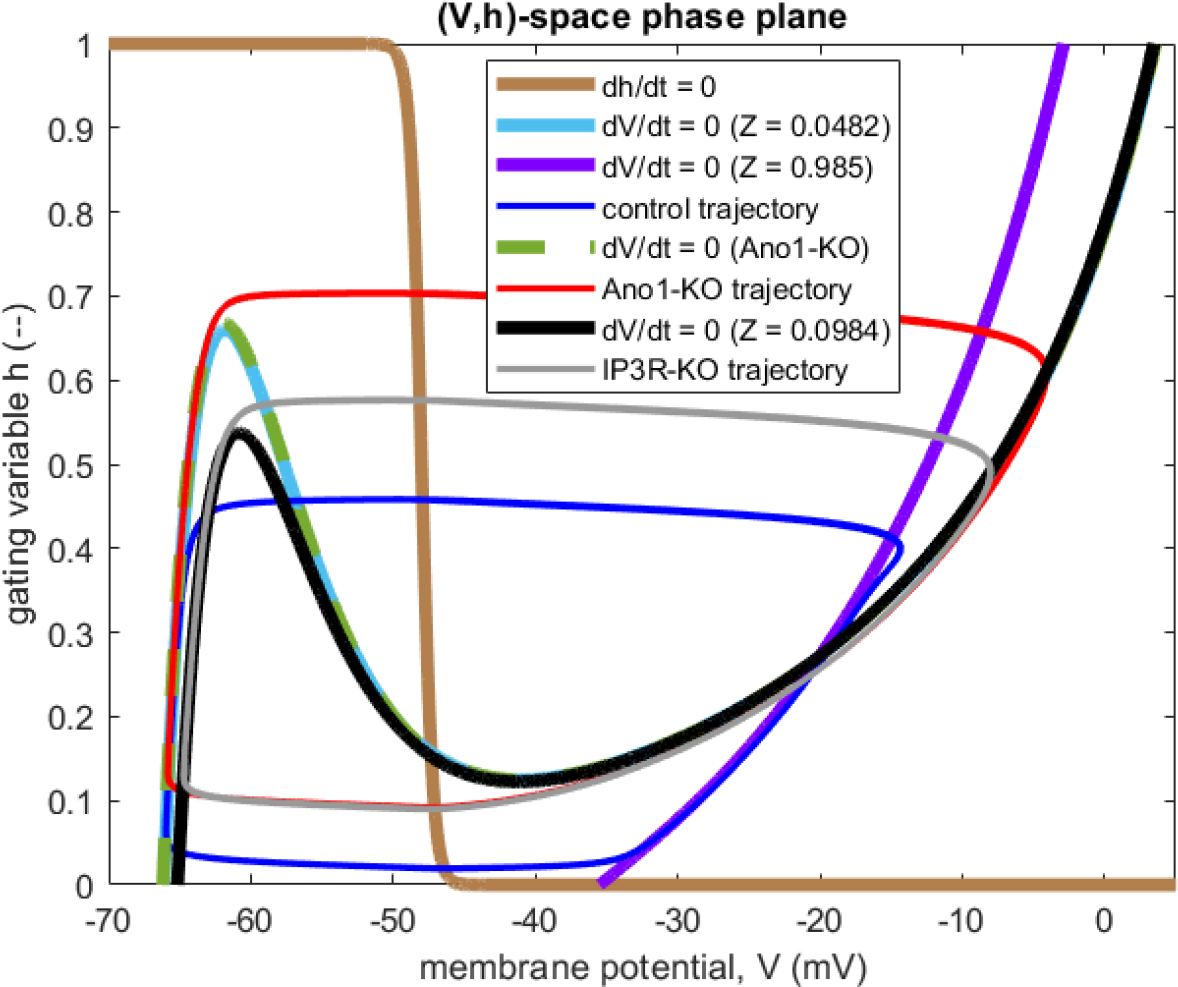
(*V*,*h*)-space phase plane for the *V*(*t*) traces shown in Fig. 6. *V*-nullclines and the unchanging *h*-nullcline are shown (thick curves), along with control (dark blue), Ano1-KO (red) and IP_3_R-KO (grey) orbital trajectories after transient decay (thin curves). Under control conditions, the *V*-nullcline moves between the extremes shown in light blue and purple. The *V*-nullcline is fixed under Ano1-KO (green) and IP_3_R-KO (black).

**Figure 8.**
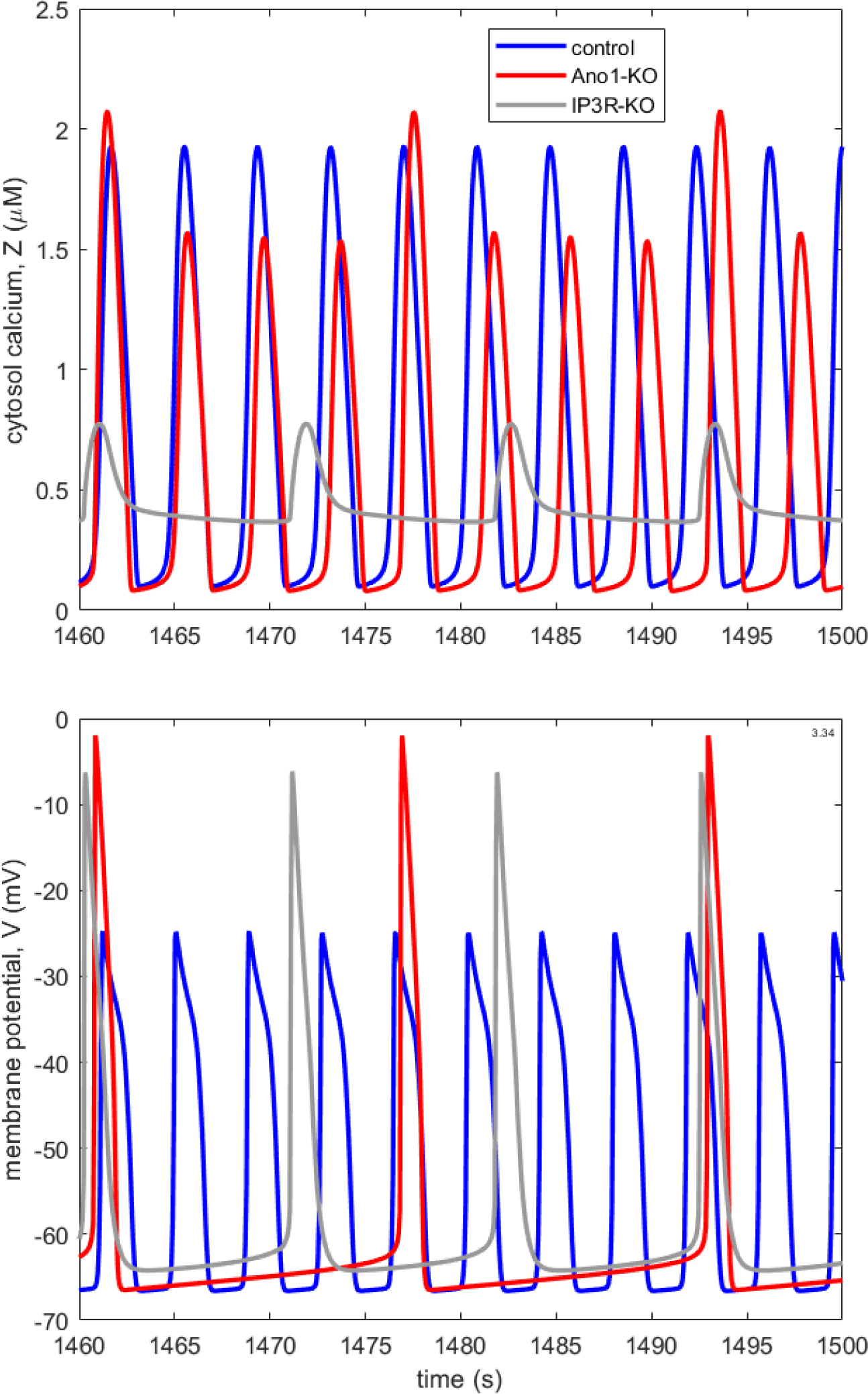
As for Fig. 6, but with *c* = 1 and parameter values as in Table 1. The AP frequencies are 15.7 (control), 3.7 (Ano1-KO) and 5.6 (IP_3_R-KO) /min.

Both IP_3_R-KO and control conditions modify the calcium oscillations from the Ano1-KO reference case (upper panel of Fig. 6). For IP_3_R-KO, the constant but non-zero [Ca^2+^] results in an increase in the frequency and decrease in amplitude of the membrane potential oscillations. In the control condition, the oscillating [Ca^2+^] also results in an increase in the frequency and decrease in amplitude of the voltage oscillations. In this case the frequency of the voltage oscillations is identical to that of the calcium oscillations (cf. the two panels of Fig. 6). In contrast, the M-clock and C-clock oscillatory frequencies are not the same in the Ano1-KO case. In the control condition, oscillatory *Z* applied to the Ano1 channel has caused the M-clock to synchronise to the oscillations in the C-clock. Thus, the action potential frequency is determined by the calcium flash frequency in the control condition, while it is determined simply by the steady [Ca^2+^] level in the IP_3_R-KO case.

Phase-plane analysis was used to investigate why this system behaviour occurs. Figure 7 shows the orbital trajectories following the left and right arms of the *V*-nullcline, as discussed in the background section. The action potential is triggered when these trajectories reach the peak of the *V*-nullcline’s left arm and jump to the right arm (the default value of *K*_h_ = 4 μM for the half-saturation [Ca^2+^] of the *h*_∞_ *Z*-dependence ensures that the *h*-nullcline is stationary). The position of this *V*-nullcline peak (and whether there is one) varies, depending on the instantaneous [Ca^2+^], *Z*(*t*). But there are two cases under which this occurs. First, in both Ano1-KO and IP_3_R-KO, the action potential occurs when the trajectory surmounts a fixed *V*-nullcline peak due to fixed *Z* (zero for Ano1-KO, non-zero for IP_3_R-KO). Constant values of *Z* greater than zero result in a lower peak, and consequently lead to higher AP frequencies and smaller AP amplitudes. But in the control condition, the action potential occurs when the trajectory surmounts a *V*-nullcline peak that is changing with time due to changing *Z*. In this case, the cyclic *Z*(*t*) increases until the *V*-nullcline peak is depressed enough for an action potential to occur. So the action potential always occurs on the upstroke of the *Z* waveform, and is thus synchronised to the C-clock oscillations.

#### 4.1.2 Coupled clocks and IP_3_R-KO data

We next return to the full model where the M-clock and the C-clock are bi-directionally coupled together. To achieve this, we set *c* = 1 in eq. 1 above so that the L-type calcium channel directly alters both the calcium concentration and the membrane potential. Figure 8 shows superimposed time-traces from model runs under IP_3_R-KO, Ano1-KO and control conditions. All three oscillations have different forms. IP_3_R-KO leads to oscillations of membrane potential that are larger in amplitude and period than control. Ano1-KO oscillations have yet larger amplitude and period. The relative order of these oscillation periods qualitatively matches the experimental data shown in Figs. 1 and 2, resolving the issue with the existing model (Hancock et al. 2022) which was discussed above.

In Fig. 8, we can study the action potential trigger by comparing the oscillations in the M-clock with those in the C-clock. Under Ano1-KO, the [Ca^2+^] oscillations, *Z*(*t*), have a different frequency^6^ to those of the membrane potential, *V*(*t*). The C-clock is not triggering oscillations in the M-clock, because the Ano1 coupling of the C-clock to the M-clock is missing. The peak of the [Ca^2+^] flash (peak *Z*) varies from one flash to the next. It is particularly high when there is an action potential, this being caused by the inrush of calcium ions via the L-type calcium channel. In contrast, under control conditions the oscillations of *V*(*t*) and *Z*(*t*) are synchronised; the calcium oscillations effect the triggering of the AP by the outrush of chloride anions via the Ano1 channel.

Under IP_3_R-KO, the oscillations of *V*(*t*) and *Z*(*t*) are again synchronised, as in the control case. However, as can be observed in the M-clock phase plane in Figure 9a, the action potential is triggered near the *V*-nullcline peak, which occurs when *Z* takes its minimum value of *Z* = 0.368 μM. To that extent, the orbital trajectory for IP_3_R-KO resembles that for Ano1-KO, and differs from that for control conditions. In the control situation, the AP is launched when *h*(*t*) is around 0.21, whereas the left-arm peak of the *V*-nullcline reaches *h* = 0.77 at another time during the cycle. Thus in control conditions rising *Z*(*t*) causes the *V*-nullcline peak to descend to meet the slowly rising operating point during the diastolic depolarization phase of the cycle. The peak descent is in this case greater than the operating-point rise, whereas under IP_3_R-KO, for this particular set of parameter values the peak descent before the AP starts is negligible.

**Figure 9.**
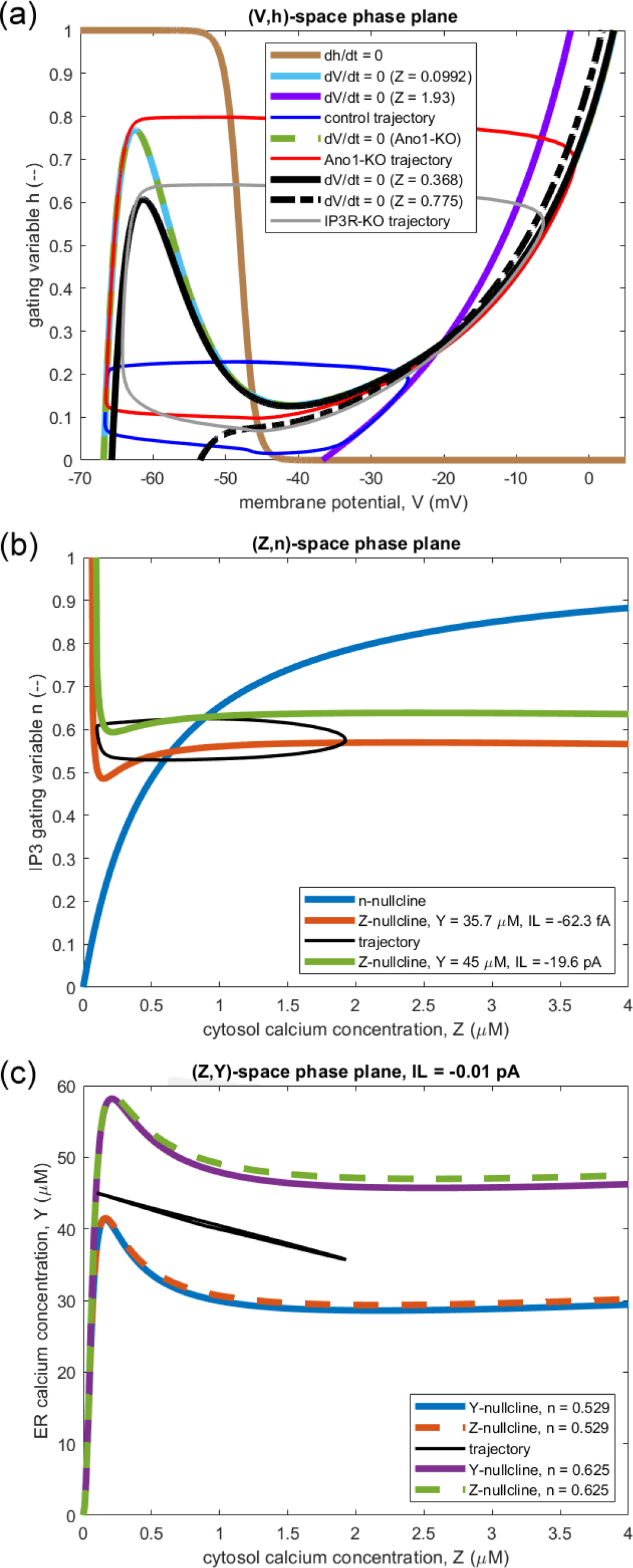
Phase planes for the three simulations shown in Fig. 8. (a): the (*V*,*h*)-space phase plane for the 2-dimensional M-clock. *V*-nullclines are shown for the minimum and maximum *Z*-values reached during a cycle of oscillation for the control conditions (thick light blue and purple curves respectively) and the IP_3_R-KO state (solid and dot-dash thick black curves). Thin curves show the orbital trajectories: dark blue = control, red = Ano1-KO, grey = IP_3_R-KO (as in Fig. 8). (b) and (c): (*Z*,*n*)-space and (*Z*,*Y*)-space phase planes for the 3-dimensional C-clock, with orbital trajectories shown in black. (b): the *Z*-nullcline is shown for the minimum *Y* and maximum *I_L_* (brown) and the maximum *Y* and minimum *I_L_* (green) reached during a cycle of oscillation^8^. (c): *I_L_* is fixed at −0.01 pA and *n* takes its cycle-extreme values.

Fig. 9b shows the (*Z*,*n*)-space phase plane^7^, including the static *n*-nullcline and approximations of the two extreme positions of the dynamic *Z*-nullcline corresponding to minimum *Y* and maximum *I_L_* in one case, and maximum *Y* and minimum *I_L_* in the other (approximate because *Y* and *I_L_* do not move exactly in antiphase). The position of the nullcline intersection varies with the *Z*-nullcline over this range. To test the stability of the intersection point, a reduced model was built in which *Y* was reduced to a linear function of *Z*, fitting the close approximation to such linear dependence shown by the trajectory in Fig. 9c. This reduces the C-clock to two dimensions, thus enabling use of the test outlined in section 3.3. For the extreme positions of the *Z*-nullcline in (*Z*,*n*)-space in the reduced model, the nullcline intersection was oscillatory but stable for the small-magnitude *I_L_*, and oscillatory and unstable for the larger-magnitude *I_L_*, showing that the reduced C-clock was capable of oscillating autonomously. Furthermore, when the full three-dimensional C-clock was isolated by setting a fixed value of *I_L_* corresponding to the mean of the values traversed cyclically when *I_L_* was set by the M-clock (−2.49 pA), trajectories diverged from the *Z*- and *n*-nullcline intersection point. The C-clock is thus operating in a region where it can drive its own oscillations. This contrasts with the trajectories from the existing model in Fig. 4b, which circle around a nullcline intersection far to the right which, by the section 3.3 test, can be shown to be a stable equilibrium point. The C-clock in that model was therefore not self-sustaining oscillation, and so the observed C-clock oscillations were being driven by the M-clock. This observation matches with the simulations in Fig. 6 showing C-clock oscillations in *Z* for Ano1-KO that are independent of, and thus not driven by, the M-clock oscillations in *V*. So, in this new model the C-clock and M-clock oscillators are coupled to and driving each other, unlike the existing-model case shown in Fig. 4, in which the M-clock was the sole driver.

Fig. 9c shows that the orbital trajectory in the (*Z*,*Y*)-space phase plane traces almost a straight line, with *Y* decreasing as *Z* increases, and *vice versa*. The slope *dY/dZ* of this line is −5.0, i.e. the magnitude is almost the same as the ratio *v*_c_/*v*_s_ = 5.4 (Table 1). This shows that, at least for the parameters used in this particular simulation, the total amount of calcium in the cell is relatively constant and is an approximately conserved quantity. The dynamics of intracellular [Ca^2+^] are thus dominated by the calcium passing between the cytosol and the endoplasmic reticulum. Mathematically, this relationship also suggests that, for the parameters used in this simulation, with *Y* essentially determined by *Z*, the (*Z*,*n*)-space phase plane is the appropriate one to analyse in order to understand the C-clock behaviour. The *Z*,*Y* plane also shows that the *Y*- and *Z*- nullclines are very similar, and the trajectories in this plane are driven by changes in the nullclines due to changes in the IP_3_R gating variable *n*.

### 4.2 Comparisons with experiment

#### 4.2.1 Variation of frequency with [IP_3_]

One of the goals of creating models of lymphatic muscle cell pace-making is to understand the mechanisms whereby the frequency of spontaneous contractions increases with vascular distending pressure [e.g., see fig. 2 of Bertram et al. (2017) for data from an isolated segment of rat mesenteric lymphatic vessel]. In lymphatic endothelial cells, a mechano-sensitive ion channel, Piezo1, mediates the sensing of oscillatory fluid shear stress (Choi et al. 2019), but Piezo1 deletion from vascular SMCs does not affect pressure sensing [see supplementary figs. 3 and 4 of Retailleau et al. (2015); we confirm the same in LMCs in yet-to-be-published work]. Thus the ion channels responsible for this pressure/frequency dependence have yet to be identified. However, variations in IP_3_ concentration in the LMC lead to variations in the frequency of pace-making in models, including this one. The [IP_3_] variations are primarily driven by the quantity of external agonist (a neurotransmitter or hormone) which binds to receptors in the cell membrane. In a cascade of subsequent reactions, G-protein and phospholipase C are activated, ultimately causing IP_3_ to be released into the cytosol (see Dupont et al. 2016 for details). However, it is known that the cell membrane G-protein-coupled receptors (see Fig. 5) are also stretch-dependent (Mederos y Schnitzler et al. 2008, Erdogmus et al. 2019), so this pathway provides one possible mechanism of pressure dependence.

In our model, cytosolic [IP_3_] has so far been taken to be constant. If it increases, more calcium ions are released from the ER, all other things being equal, and the C-clock is thus impelled to speed up. Assuming the pressure dependence of the G-protein-coupled receptors, we can therefore use cytosolic [IP_3_] as a surrogate for distending pressure. Figure 10 shows how the frequency of APs and the associated calcium flashes varies with cytosolic [IP_3_] in our model. The figure also shows equivalent data for the model published by Hancock et al. (2022), and for the resynthesised version of the model by Imtiaz et al. (2007) that was implemented for that paper.

**Figure 10.**
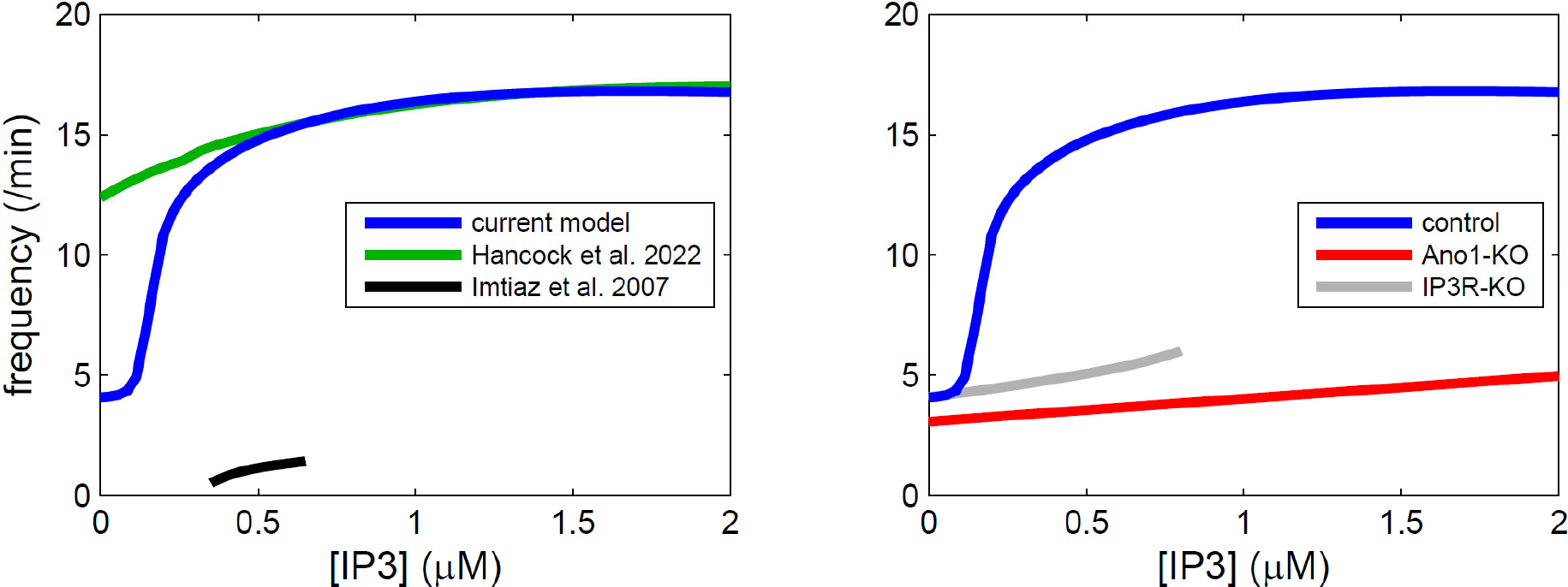
Pacemaker frequency as a function of IP_3_ concentration. Curves end where the model in question no longer oscillated without changing other parameters. Left panel: black, the model of Imtiaz et al. (2007) as reinterpreted by Hancock et al. (2022). Green, the model of Hancock et al. (2022). Blue, the new model. Right panel: the new model under control (blue), Ano1-KO (red) and IP_3_R-KO (grey).

It is seen (left panel) that the frequency range of IP_3_ concentrations and frequencies over which the reinterpreted Imtiaz model provided pacemaker oscillations was very limited^9^. Pace-making over a much wider range of [IP_3_] was provided by the model of Hancock et al. (2022), but the range of pacemaker frequencies was still limited. In contrast, under control conditions as for Figs. 8 and 9, the present model ranges between a minimum frequency of 4.1 /min at *P* = 0 μM, and a maximum of 16.8 /min at *P* = 2 μM. For comparison, the *ex vivo* data of Bertram et al. (2017) display a yet wider frequency range, from less than 2 /min at zero transmural pressure, to some 28 /min at 50 cmH_2_O. But the frequency-[IP_3_] pattern shown here, saturating at high [IP_3_], is suggestively similar to the frequency-pressure pattern found experimentally, saturating at high distending pressure.

The right panel shows that, for all IP_3_ concentrations from 0.2 to 2 μM, in the new model, Ano1-KO reduces the AP frequency to between 24 and 30% of the corresponding frequency under control conditions. This finding exactly parallels what we found experimentally in mouse lymphatic collecting vessels (Zawieja et al. 2019): deleting Ano1 using three different smooth-muscle-specific Cre lines, a similar result was found in each, i.e. the basal pace-making rate was reduced to <25% of control and pressure-induced chronotropy was blunted or abolished.

The right panel also shows how AP frequency is reduced in IP_3_R-KO in the new model: for all IP_3_ concentrations from 0.2 to 0.8 μM (beyond which IP_3_R-KO oscillation ceases unless other parameters are adjusted), AP frequency is reduced to between 34 and 41% of control. At [IP_3_] = 0, the control and IP_3_R-KO traces become one. This matches fairly closely the experimental results (Zawieja et al., submitted), which show that contraction frequency in IP_3_R1-KO vessels was between 17% and 27% of IP_3_R1 fl/fl control counterparts across the pressure range that was studied. Specifically, at the distending pressure of 3 cmH_2_O, IP_3_R1-KO frequency was 26.9% of control (4.45 vs. 16.53 /min).

#### 4.2.2 Calcium dependence of L-type current

In all the new-model simulations shown thus far, the *Z*-dependence of the gating variable *h*(*V*,*Z*) has been suppressed for the sake of simplicity, by setting a value for *K*_h_ not reached during the *Z* oscillation. Finally, this dependence is now reintroduced, by setting *K*_h_ = 0.6 μM. Figure 11 shows how this change affects the waveforms of *Z*(*t*) and *V*(*t*), and the M-clock phase plane (the C-clock phase planes are essentially unchanged from Fig. 9).

**Figure 11.**
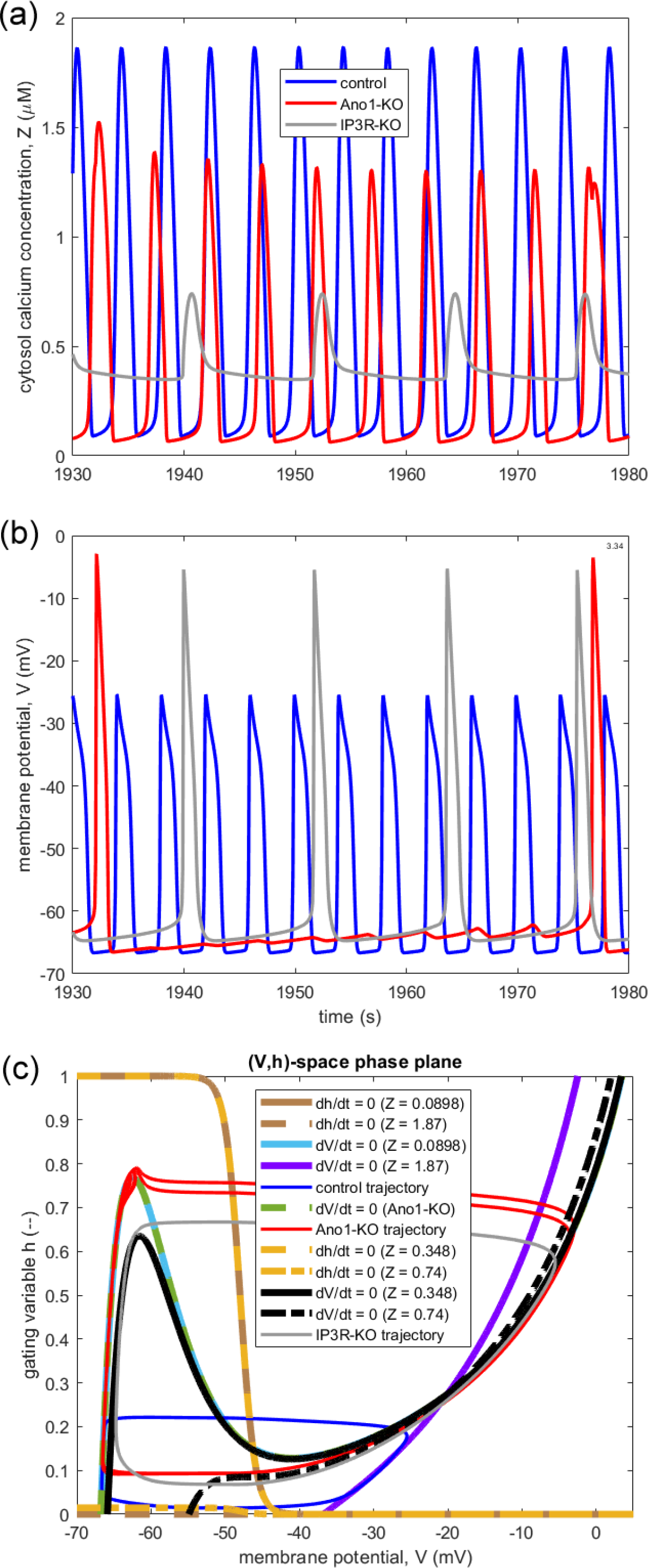
(a) and (b): waveforms as in Fig. 8, but with *K*_h_ = 0.6 μM. (c): the corresponding (*V*,*h*)-space phase plane. The *h*-nullcline is shown for the minimum and maximum of the control *Z*-waveform (brown: solid and dashed respectively), and for the minimum and maximum of the IP_3_R-KO *Z*-waveform (yellow: dashed and dash-dot respectively). See legend for other (*V*,*h*)-space trace identifiers.

The effects are especially marked during Ano1-KO. Insofar as *Z*(*t*) is concerned, the lack of cyclic regularity, earlier footnoted, is now highly evident. With Ano1-KO removing clock coupling, so that the M-clock frequency is untied from that of the C-clock, the two clocks oscillate at incommensurate rates. However, the size of the [Ca^2+^] flash is affected by APs, but to an extent which depends on the timing of the AP relative to the flash. At the beginning of the time-window shown (long after numerical start-up transients have decayed), the AP occurs at the same time as the *Z* flash, causing the *Z* peak to be higher than usual. Over the subsequent few *Z* flashes, the peak height decays back to its unperturbed value. But the next AP occurs slightly later than the corresponding *Z* flash, the peak of which is accordingly barely higher than the preceding one. But the AP causes a late rebound in [Ca^2+^], so that the overall effect is to lengthen the flash.

With the reactivation of the *Z*-dependence of the L-type gate function *h*, the effects of the C-clock oscillation are also evident in the membrane potential trace for Ano1-KO, i.e. there is C- to M-clock coupling. The frequency of Ano1-KO APs is now even further reduced relative to both control conditions and the corresponding frequency seen in Fig. 8. As with the time-course of intracellular [Ca^2+^], the interaction of the two clocks via oscillations of incommensurate frequency means that there is not a strictly repetitive *V*(*t*) waveform, but a calculation based on the timing of the two Ano1-KO AP peaks shown yields a pseudo-frequency of 1.33 /min. The corresponding frequency for the (properly repetitive) Ano1-KO APs in Fig. 8 is 3.74 /min.

Of even greater interest is the fact that, during the diastolic slow depolarisation phase, the membrane potential trace for Ano1-KO in Fig. 11 now shows perturbations arising from the [Ca^2+^] oscillations. In other words, the operation of the C-clock can here be observed via the M-clock’s output. This is a direct result of the effect of changing *Z* on the *h*-nullcline (bottom panel of Fig. 11), the left arm of which now moves during the cycle between *h* = 1 and *h* = 0.015. This opens the possibility of observing the same thing experimentally. Figure 12 shows experimental data of membrane potential which include a qualitatively similar phenomenon occurring during the slow depolarisation phase of Ano1-KO, here induced by administration of the pharmacological agent Ani9 to inhibit Ano1 instead of by genetic knock-out. As in the numerical trace of Fig. 11, diastolic oscillatory perturbations of increasing amplitude occur. While it is not possible to corroborate this speculation in the absence of a simultaneous recording of [Ca^2+^] flash frequency (not technically possible in our laboratory at this time), it is possible that the frequency of these otherwise unexplained experimental oscillations may represent the operation of the real C-clock in this case. If so, then as in the numerical simulation, the C-clock frequency in Ano1-KO greatly exceeds that of the M-clock, and can be approximated from the trace as 11.5 /min. The frequency of the lesser Ano1-KO *Z* peaks (between the two that are affected by APs) in the top panel of Fig. 11 is 12.0 /min.

**Figure 12.**
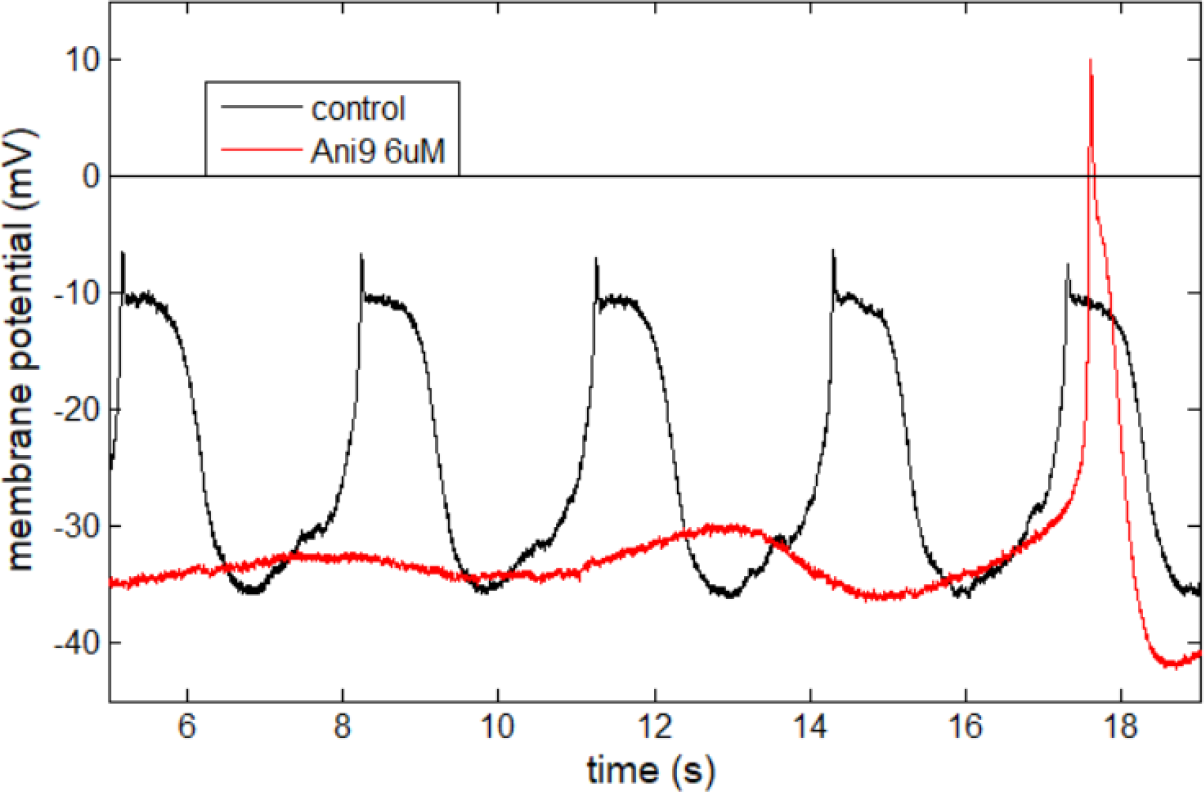
Recording of action potentials from an isolated mouse inguinal-axillary lymphatic vessel under control conditions (black) and after administration of 6 μM Ani9 to induce Ano1 channel block (red).

## 5 DISCUSSION

In this work, our overarching goal is to understand the key mechanisms and system effects that drive pace-making in lymphatic muscle cells. To achieve this goal, we have focused on determining the underlying oscillators in lymphatic muscle cells and how they are coupled. The focus of this interest has been on the M-clock and C-clock; determining which of the clocks is driving oscillations and how the clocks are coupled. An important part of this determination is studying how individual mechanisms such as the IP_3_ receptor affect the individual clocks and the system as a whole.

We have made a number of changes to our existing LMC model (Hancock et al. 2022), which focus on modifying the mechanisms in the C-clock model. These changes had two motivations. First, the experimental data used in this paper involve the knock-out of a key C-clock mechanism (IP_3_R), and so developing an accurate model of the C-clock was important for these data. Second, our existing model was based on a model by Imtiaz et al. (2007) which added dependent M-clock components to an existing C-clock model by Dupont & Goldbeter (1993). The Dupont-Goldbeter model relied on store depletion to reduce IP_3_R current, contrasting with more recent models that incorporate late IP_3_R inhibition. The latter is now viewed as a crucial feature of calcium oscillations and so the Dupont-Goldbeter model is no longer widely used (Dupont et al. 2016). As such, the IP_3_R mechanism in the model (DeYoung & Keizer 1992) was brought in line with more recent C-clock models in other cell types. The result of the IP_3_R channel modification is a model structure that has similarities to the existing model, but with an extra gating variable for the IP_3_ receptor. This addition takes the number of variables required for the C-clock model to three^10^. Apart from this change, the ion channels represented are equivalent to the existing model, with only minor modifications to kinetics. However, the added complexity to the IP_3_R model can dramatically change the behaviour of the C-clock. In particular, analysis of the self-driven oscillations in the C-clock is centred on the intracellular calcium concentration *Z* and IP_3_R gate state *n*, rather than *Z* and the intra-store calcium concentration *Y* as in the existing model. This can be observed in the phase planes, where C-clock oscillations are best viewed in the (*Z*,*n*)-space phase plane rather than in (*Z*,*Y*)-space (see Fig. 9).

The proposed model is a system with dual coupled clocks, where both are important for initiation of the action potential. This structure results in the two clocks synchronising in frequency. Dual-clock coupled behaviour has been found in cardiac muscle cells, via both modelling and experimental results (Maltsev & Lakatta 2009, 2013), but has not previously been found in LMCs. The modelling for cardiac cells involves highly detailed mechanistic models with large numbers of equations; see, e.g. DiFrancesco & Noble (1985). In contrast, the model presented here is relatively low order, while obtaining qualitatively similar results. This lower order enables us gain insight from analysis that would not be possible with a more complex model. In particular, it enabled the use of phase-plane analysis to understand the coupling between the two clocks.

Despite the change to the C-clock model, the calcium concentration *Z* can still be used to represent the C- to M-clock coupling, which was an important property of our existing model (Hancock et al. 2022). Similarly, the L-type current *I_L_* can still best represent the M- to C-clock coupling. These properties enable each coupling direction to be represented by a single variable, simplifying the models and enabling insights to be drawn from the analysis. The alternative is to use *h* and *V* together to represent the coupling from the M- to the C-clock, which is unnecessarily complex and obscures the real mechanism.

We have demonstrated that coupling can occur between the clocks by multiple different means. The results showed that the C-clock can affect the oscillations in the M-clock either via a constant level of calcium concentration *Z* setting the height of a static V-nullcline threshold for the operating point to surmount, or via oscillations in the calcium concentration moving the action-potential threshold dynamically. Both methods act via the Ano1 mechanism, but their system effects are different. In the first case, the higher the steady-state *Z*, the lower the level of the gating variable *h* at which the M-clock self-triggers an action potential. In this case C-clock oscillations are not required for action potentials. In the second case, the rising phase of the cyclic *Z* oscillation progressively reduces the M-clock’s *h*-level trigger point for the action potential. If this point falls below the current state of *h* then an action potential is initiated. Thus the C-clock effectively triggers the M-clock’s action potential, rather than changing the self-trigger. The upswing in oscillating calcium concentration then necessarily coincides with the triggering of the action potential in the M-clock, synchronising the two clocks.

The second of these cases represents a situation which goes beyond the traditional physiological view of the action potential happening at such time as slow diastolic depolarisation causes the membrane potential to surpass a static voltage threshold. Instead we show that, whenever calcium flashes and action potentials are synchronised, the barrier or threshold itself is dynamically falling away at the instant when the action potential is triggered. Furthermore, we see that, in both cases, the M-clock threshold is far more effectively viewed as a level of the L-type gating variable *h*, rather than as a level of the membrane potential *V*. Obviously, for the experimentalist, the only measurable quantity remains the membrane potential^11^, and *V*(*t*) and *h*(*t*) are indeed linked monotonically albeit nonlinearly during the cycle phase leading up to the next action potential. But, expressed in terms of membrane potential, these thresholds and threshold motions are a small fraction of the total excursion, whereas in terms of gating variable *h* they are order-1 quantities. There is also coupling from the C-clock to the M-clock via inactivation of the L-type calcium channel, but this mechanism has no effect on the triggering in this model.

The M-clock coupling to the C-clock occurs via a large inrush of calcium ions during the action potential via the L-type calcium channel. If the C-clock is not oscillating independently then this periodic inrushing current causes the C-clock to oscillate passively in response. Alternatively, if the C-clock oscillations trigger an action potential in the M-clock then the inrush of current amplifies the increase in cytosolic calcium concentration. This amplification causes a [Ca^2+^] flash of larger amplitude and longer duration in the C-clock. Overall, the C-clock can initiate an earlier action potential in the M-clock, while the M-clock causes amplification of the calcium uptake. Combined, this bidirectional coupling can cause the two to synchronise in frequency.

## 6 CONCLUSION

Modelling is an important tool to help understand pace-making in lymphatic muscle cells, and consequently it is important for understanding lymphatic system dysfunction. In this paper we have extended the state-of-the-art in pacemaker modelling for lymphatic muscle cells. We have introduced a model of LMCs where the M-clock and C-clock are synchronised and both driving the action potentials and associated calcium concentration surges. This change resulted from changes to the mechanism of the IP_3_R channel in the C-clock, and enables the model to match experimental data for IP_3_R knock-out. We have also provided phase-plane analysis explaining the result. The model, results and analysis are important for progressing comprehension of pace-making and its dysfunction in lymphatic muscle cells.

## ACKNOWLEDGEMENTS

This research, EJH, and in part CDB, were supported by NIH grant R01-HL-122578 to MJD. SDZ acknowledges NIH grant R00-HL-143198.

## Footnotes

1 Arguably a similar degree of complexity is manifested by the smooth muscle in the wall of those small arterial blood vessels which exhibit vasomotion on a rather rapid time-scale, in addition to their traditionally recognised function of sustaining a standing level of vascular tone and thus hydraulic resistance.

2 In this paper we term the calcium-concentration equivalent of the action potential a calcium flash.

3 There are three IP_3_ receptor isoforms; it has not yet been ruled out that there could be a degree of compensatory expression of IP_3_R2 or IP_3_R3.

4 However, these ratios are not particularly meaningful, since the control and IP_3_R-KO traces come from different animals.

5 We here follow precedent (Dupont et al. 2016) in assuming that the plasma membrane Ca^2+^ ATPase pump operation is stoichiometrically neutral (as many positive charges on monovalent hydrogen ions are imported as are exported on bivalent calcium ions), and thus no term in *J*_ce_ appears in eq. 4. The exact stoichiometry is uncertain (Brini & Carafoli 2011).

6 In fact there is no true frequency to the *Z*(*t*) oscillations under Ano1-KO, because the underlying C-clock frequency is not synchronised with the APs, even at the apparent reduced rate of one AP per four flashes—there is continuous slippage of the AP phase relative to the latest flash. In consequence the amount to which L-type current contributes to the largest flashes varies over a longer passage of time than is shown here. In dynamical systems terms, the oscillation is quasi-periodic.

7 In fact, this is rather a projection onto two dimensions of the three-dimensional (*Z*,*Y*,*n*)-space, since the trajectory does not keep to a constant value of *Y*. Similarly, the two positions shown for the *Z*-nullcline are for different values of Y.

8 The usual convention is observed here that positive current entails the exit of positive ions; thus the L-type current, involving influx of positive calcium ions, is negative. Therefore the minimum value corresponds to the greatest magnitude.

9 In their fig. 9, Imtiaz et al. (2007) demonstrated a wider [IP3] range for oscillation in the Dupont-Goldbeter C-clock-only model upon which their model was founded, but we here refer to their entire model as reinterpreted on a more physical basis and with more realistic parameter values by Hancock et al. (2022).

10 Much more complex models for the IP_3_ receptor exist, but these necessitate many ODEs, taking the resulting ODE system out of the realm where even approximate phase-plane analysis is feasible or useful.

11 Experimentally quantifying the L-type current activation and inactivation range in LMCs would help increase the accuracy with which the model can fit the real situation.

## REFERENCES

Adams KE, Rasmussen JC, Darne C, Tan I-C, Aldrich MB, Marshall MV, Fife CE, Maus EA, Smith LA, Guilloid R, Hoy S, Sevick-Muraca EM (2010) Direct evidence of lymphatic function improvement after advanced pneumatic compression device treatment of lymphedema. Biomedical Optics Express 1(1): 114–125.

Bertram CD, Macaskill C, Davis MJ, Moore JR. JE (2017) Valve-related modes of pump failure in collecting lymphatics: numerical and experimental investigation. Biomechanics and Modeling in Mechanobiology 16(6): 1987–2003. doi:10.1007/s10237-017-0933-3

Breslin JW, Yang Y, Scallan JP, Sweat RS, Adderley SP, Murfee WL (2019) Lymphatic vessel network structure and physiology. Comprehensive Physiology 9: 207–299. doi:10.1002/cphy.c180015

Brini M, Carafoli E (2011) The plasma membrane Ca^2+^ ATPase and the plasma membrane sodium calcium exchanger cooperate in the regulation of cell calcium. Cold Spring Harbor Perspectives in Biology 3(2): a004168 (15 pp.). doi:10.1101/cshperspect.a004168

Choi D, Park E, Jung E, Cha B, Lee S, Yu J, Kim PM, Lee S, Hong YJ, Koh CJ, Cho C-W, Wu Y, Jeon NL, Wong AK, Shin L, Kumar SR, Bermejo-Moreno I, Srinivasan RS, Cho I-T, Hong Y-K (2019) Piezo1 incorporates mechanical force signals into the genetic program that governs lymphatic valve development and maintenance. JCI Insight 4(5): e125068 (15 pp.). doi:10.1172/jci.insight.125068

De Young GW, Keizer J (1992) A single-pool inositol 1,4,5-trisphosphate-receptor-based model for agonist-stimulated oscillations in Ca^2+^ concentration. Proceedings of the National Academy of Sciences of the USA 89(20): 9895–9899. doi:10.1073/pnas.89.20.9895

DiFrancesco D, Noble D (1985) A model of cardiac electrical activity incorporating ionic pumps and concentration changes. Philosophical Transactions of the Royal Society B 307(1133): 353–398. doi:10.1098/rstb.1985.0001

Dupont G, Goldbeter A (1993) One-pool model for Ca^2+^ oscillations involving Ca^2+^ and inositol 1,4,5-trisphosphate as co-agonists for Ca^2+^ release. Cell Calcium 14(4): 311–322

Dupont G, Falcke M, Kirk V, Sneyd J (2016) Models of Calcium Signalling. Interdisciplinary Applied Mathematics, vol 43. Springer International Publishing, Switzerland. doi:10.1007/978-3-319-29647-0

Emrich SM, Yoast RE, Xin P, Arige V, Wagner LE, Hempel N, Gill DL, Sneyd J, Yule DI, Trebak M (2021) Omnitemporal choreographies of all five STIM/Orai and IP_3_Rs underlie the complexity of mammalian Ca^2+^ signaling. Cell Reports 34: 108760 (13 pp. + 6 pp.). doi:10.1016/j.celrep.2021.108760

Erdogmus S, Storch U, Danner L, Becker J, Winter M, Ziegler N, Wirth A, Offermanns S, Hoffmann C, Gudermann T, Mederos Y Schnitzler M (2019) Helix 8 is the essential structural motif of mechanosensitive GPCRs. Nature Communications 10(1): 5784 (15 pp.). doi:10.1038/s41467-019-13722-0

Goldbeter A, Dupont G, Berridge MJ (1990) Minimal model for signal-induced Ca^2+^ oscillations and for their frequency encoding through protein phosphorylation. Proceedings of the National Academy of Sciences of the USA 87(4): 1461–1465. doi:10.1073/pnas.87.4.1461

Hald BO, Castorena-Gonzalez JA, Zawieja SD, Gui P, Davis MJ (2018) Electrical communication in lymphangions. Biophysical Journal 115(5): 936–949. doi:10.1016/j.bpj.2018.07.033

Hancock EJ, Zawieja SD, Macaskill C, Davis MJ, Bertram CD (2022) Modelling the coupling of the M-clock and C-clock in lymphatic muscle cells. Computers in Biology and Medicine 142: 105189-1–105189-12. doi:10.1016/j.compbiomed.2021.105189

Hille B (2001) Ion Channels of Excitable Membranes, 3rd edn. Sinauer Associates Inc., Sunderland MA

Hodgkin AL, Huxley AF (1952) A quantitative description of membrane current and its application to conduction and excitation in nerve. Journal of Physiology 117: 500–544

Imtiaz MS, Zhao J, Hosaka K, von der Weid P-Y, Crowe MJ, van Helden DF (2007) Pacemaking through Ca^2+^ stores interacting as coupled oscillators via membrane depolarization. Biophysical Journal 92(11): 3843–3861. doi:10.1529/biophysj.106.095687

Jørgensen MG, Toyserkani NM, Hansen FG, Bygum A, Sørensen JA (2021) The impact of lymphedema on health-related quality of life up to 10 years after breast cancer treatment. NPJ Breast Cancer 7 (1): 70 (8 pp.). doi:10.1038/s41523-021-00276-y

Keener J, Sneyd J (2009) Mathematical Physiology I: Cellular Physiology. Interdisciplinary Applied Mathematics, vol 8/I, 2nd. edn. Springer Science+Business Media LLC, New York. doi:10.1007/978-0-387-75847-3

Lee Y, Zawieja SD, Muthuchamy M (2022) Lymphatic collecting vessel: new perspectives on mechanisms of contractile regulation and potential lymphatic contractile pathways to target in obesity and metabolic diseases. Frontiers in Pharmacology 13: 848088 (19 pp.). doi:10.3389/fphar.2022.848088

Lees-Green R, Gibbons SJ, Farrugia G, Sneyd J, Cheng LK (2014) Computational modeling of anoctamin 1 calcium-activated chloride channels as pacemaker channels in interstitial cells of Cajal. American Journal of Physiology – Gastrointestinal and Liver Physiology 306(8): G711–G727. doi:10.1152/ajpgi.00449.2013

Li Y-X, Rinzel J (1994) Equations for InsP_3_ receptor-mediated [Ca^2+^]_i_ oscillations derived from a detailed kinetic model: a Hodgkin-Huxley like formalism. Journal of Theoretical Biology 166(4): 461–473

Maltsev VA, Lakatta EG (2009) Synergism of coupled subsarcolemmal Ca^2+^ clocks and sarcolemmal voltage clocks confers robust and flexible pacemaker function in a novel pacemaker cell model. American Journal of Physiology – Heart and Circulatory Physiology 296(3): H594–H615. doi:10.1152/ajpheart.01118.2008

Maltsev VA, Lakatta EG (2013) Numerical models based on a minimal set of sarcolemmal electrogenic proteins and an intracellular Ca^2+^ clock generate robust, flexible, and energy-efficient cardiac pacemaking. Journal of Molecular and Cellular Cardiology 59: 181–195. doi:10.1016/j.yjmcc.2013.03.004

McAllister RE, Noble D, Tsien RW (1975) Reconstruction of the electrical activity of cardiac Purkinje fibres. Journal of Physiology 251: 1–59

Mederos y Schnitzler M, Storch U, Meibers S, Nurwakagari P, Breit A, Essin K, Gollasch M, Gudermann T (2008) Gq-coupled receptors as mechanosensors mediating myogenic vasoconstriction. EMBO Journal 27(23): 3092–3103. doi:10.1038/emboj.2008.233

Moore JR. JE, Bertram CD (2018) Lymphatic system flows. Annual Review of Fluid Mechanics 50: 459–482. doi:10.1146/annurev-fluid-122316-045259

Morris C, Lecar H (1981) Voltage oscillations in the barnacle giant muscle fiber. Biophysical Journal 35: 193–213

Mortimer PS, Rockson SG (2014) New developments in clinical aspects of lymphatic disease. Journal of Clinical Investigation 124(3): 915–921. doi:10.1172/JCI71608

Oliver G, Kipnis J, Randolph GJ, Harvey NL (2020) Review - The lymphatic vasculature in the 21st century: novel functional roles in homeostasis and disease. Cell 182(2): 270–296. doi:10.1016/j.cell.2020.06.039

Olszewski WL, Engeset A (1980) Intrinsic contractility of prenodal lymph vessels and lymph flow in human leg. American Journal of Physiology – Heart and Circulatory Physiology 239(6): H775–H783

Retailleau K, Duprat F, Arhatte M, Ranade SS, Peyronnet R, Martins JR, Jodar M, Moro C, Offermanns S, Feng Y, Demolombe S, Patel A, Honoré E (2015) Piezo1 in smooth muscle cells is involved in hypertension-dependent arterial remodeling. Cell Reports 13(6): 1161–1171. doi:10.1016/j.celrep.2015.09.072

Rockson SG (2001) Lymphedema. American Journal of Medicine 110(4): 288–295 doi:10.1016/s0002-9343(00)00727-0

Rockson SG, Rivera KK (2008) Estimating the population burden of lymphedema. Annals of the New York Academy of Science 1131: 147–154. doi:10.1196/annals.1413.014

Strogatz SH (2015) Nonlinear Dynamics and Chaos – With Applications to Physics, Biology, Chemistry, and Engineering, 2nd edn. CRC Press, Boca Raton FL

Yaniv Y, Lakatta EG, Maltsev VA (2015) From two competing oscillators to one coupled-clock pacemaker cell system. Frontiers in Physiology 6: 28 (8 pp.). doi:10.3389/fphys.2015.00028

Zawieja SD, Castorena JA, Gui P, Li M, Bulley SA, Jaggar JH, Rock JR, Davis MJ (2019) Ano1 mediates pressure-sensitive contraction frequency changes in mouse lymphatic collecting vessels. Journal of General Physiology 151(4): 532–554. doi:10.1085/jgp.201812294

Zawieja SD, Davis MJ et al., (submitted) Lymphatic muscle IP_3_R1 is required for normal murine lymphatic collecting vessels pressure-dependent chronotropy and myogenic activity.

Zhang L, Zhang H, Zhong Q, Luo Q, Gong N, Zhang Y, Qin H, Zhang H (2021) Predictors of quality of life in patients with breast cancer-related lymphedema: effect of age, lymphedema severity, and anxiety. Lymphatic Research and Biology 19(6): 573–579. doi:10.1089/lrb.2020.0073

